# Baicalin ameliorates behavioral and physiological deficits in *Gtf2i*-duplication mice and modulates synaptic activity in 7q11.23 Duplication Syndrome patient-derived neurons

**DOI:** 10.64898/2026.02.12.705480

**Authors:** David Peles, Shai Netser, Natali Ray, Taghreed Suliman, Shani Stern, Shlomo Wagner

## Abstract

7q11.23 Duplication Syndrome (7Dup) is a type of syndromic autism spectrum disorder caused by duplication of a typically 1.5-1.8 Mb segment in section q11.23 of chromosome 7, including 25-27 genes. Previous work has highlighted the *GTF2I* gene as playing a major role in the phenotype of 7Dup patients. Accordingly, mice with *Gtf2i* duplication (*Gtf2i*^+/dup^) are commonly used as an animal model of 7Dup. We previously reported deficits in several behavioral and physiological modalities, which were associated with *Gtf2i* dosage in mice conducting a battery of social discrimination tests. Here, we report the effect of treating *Gtf2i*^+/dup^ mice with Baicalin, a naturally occurring flavonoid added to the mice’s drinking water (0.15 mg/ml), on these deficits. We found that Baicalin treatment ameliorated the higher surface temperature observed in *Gtf2i*^+/dup^ males and the lower tail temperature observed in *Gtf2i*^+/dup^ females during the social behavior tests. It also prevented the increased defecation rate exhibited by *Gtf2i*^+/dup^ mice during the social preference test. We further analyzed the effect of Baicalin treatment on cortical neurons differentiated from 7Dup patient-derived IPSCs. Using whole cell patch clamp and calcium imaging, we found an increased rate of excitatory postsynaptic currents in Baicalin-treated cells, without a change in their firing rate, indicating a stronger synaptic activity in the Baicalin-treated cells. Altogether, our results reveal that Baicalin administration alleviates some of the behavioral and physiological effects of *Gtf2i* duplication in mice, and affects neuronal activity in cultured 7Dup human neurons. Thus, Baicalin administration has the potential to serve as a treatment for 7Dup patients.

## INTRODUCTION

7q11.23 Duplication Syndrome (7Dup, for short) is caused by duplication of 1.5-1.8 Mb segment in human chromosome 7. 7Dup patients are diagnosed with social anxiety, selective mutism, autism spectrum disorder (ASD), oppositional behavior, and intellectual disability in some of the patients [1, 2]. *GTF2I*, one of the 25-27 genes typically included in the duplicated segment, is considered to play a major role in the 7Dup phenotype, as suggested by rare cases with small duplications or deletions that include the *GTF2I* region [3–7]. Mice with duplication of the *Gtf2i* gene (Dup mice, for short) are commonly used as a mouse model for 7Dup [8–11]. In a previous work, we have reported multiple behavioral and physiological deficits associated with either duplication or deletion of *Gtf2i* in mice [8].

Baicalin is a naturally occurring flavonoid usually derived from the root of the *Scutellaria baicalensis*, a plant commonly used in traditional Chinese medicine [12]. Baicalin’s pharmacological effect was studied in cellular and rodent models (see [13–16] for a review), and to a limited extent also in a clinical trial in humans [17]. When administered orally, Baicalin absorption was shown to be largely mediated by the intestinal bacteria, which transform Baicalin to Baicalein, which, after absorption, transforms back to Baicalin [18].

Inhibition of the histone modification eraser enzyme lysine demethylase 1 (LSD1) and of histone deacetylase (HDAC) enzymes were previously suggested as potential treatments for 7Dup [11, 19, 20]. Our interest in Baicalin, which exerts the ability to pass the blood-brain barrier (BBB) [21, 22], stems from its LSD1 (KDM1A) inhibition properties [23, 24], as well as from the HDAC inhibition and downregulation properties of Baicalin and Baicalein [25–27]. Thus, we hypothesized that Baicalin administration may alleviate the deficits exhibited by Dup mice.

In this work, we treated Dup mice with Baicalin, dissolved in their drinking water (0.15 mg/ml) from the day of weaning to the end of the experiments. We analyzed the effect of this treatment on the behavioral and physiological variables previously found by us to be modified in Dup mice while the animals conducted a battery of social discrimination tests. In addition, we treated cortical neurons differentiated from 7Dup patient-derived IPSCs with Baicalin (5ug/ml, which is 11.2uM) and examined the effects of this treatment on the neuronal and synaptic activity using whole cell patch clamp and calcium imaging. Our results reveal that Baicalin indeed alleviates the deficits exhibited by Dup mice and affects synaptic activity in the 7Dup neuronal cultures. Altogether, these results support the potential use of Baicalin as a treatment for 7Dup.

## MATERIALS AND METHODS

### Animals

Adult subject mice (12-14 weeks old) of CD1 (ICR) genetic background were derived from breeding couples of *Gtf2i*^+/dup^ mice [9] (Dup mice, for short) and WT mice. Subject mice were bred and raised in the SPF mouse facility of the University of Haifa. Stimuli mice were adult CD1 mice purchased from Envigo (Rehovot, Israel) or raised at the University of Haifa facility. Mice were housed in groups of 3-5 with *ad libitum* food and water, in a 12 hours dark/12 hours light cycle. Experiments were performed in the dark phase of the dark/light cycle. All experiments were approved by the University of Haifa ethics committee (Reference #: UoH-IL2301-103-4) and included veterinary supervision.

### Baicalin treatment and experiment timeline in mice

Baicalin powder (Molecular Formula: C21H18O11, CAS No. 21967-41-9, purity >95%) was purchased from AvaChem Scientific, San Antonio, TX, USA (Catalog No. 1026). Mice home cages, each including both Dup mice and WT mice, were assigned as either Baicalin-treatment or Control. In the Baicalin treatment cages, Baicalin was dissolved in the mice’s drinking water (0.15mg/ml) from the day of weaning and until the end of the social behavior tests, which were done at the age of 2-5 months (Fig. 1A). Though baicalin solubility in pure water is considered very low (0.052 mg/ml according to [28]), we received a visually clear and uniform yellowish liquid when dissolving Baicalin in this concentration in tap water (which naturally contains some amount of salts), and hence we did not use additional solvents. The control cages received regular tap water. We replaced the Baicalin-dissolved drinking water in the Baicalin-treated cages three times a week. The average consumption of Baicalin dissolved water in adult mice was ∼5 ml per day per adult mouse. Assuming a typical weight of 30-35g will result in an estimated consumption of ∼21-25 mg/kg Baicalin per day.

**Figure 1:**
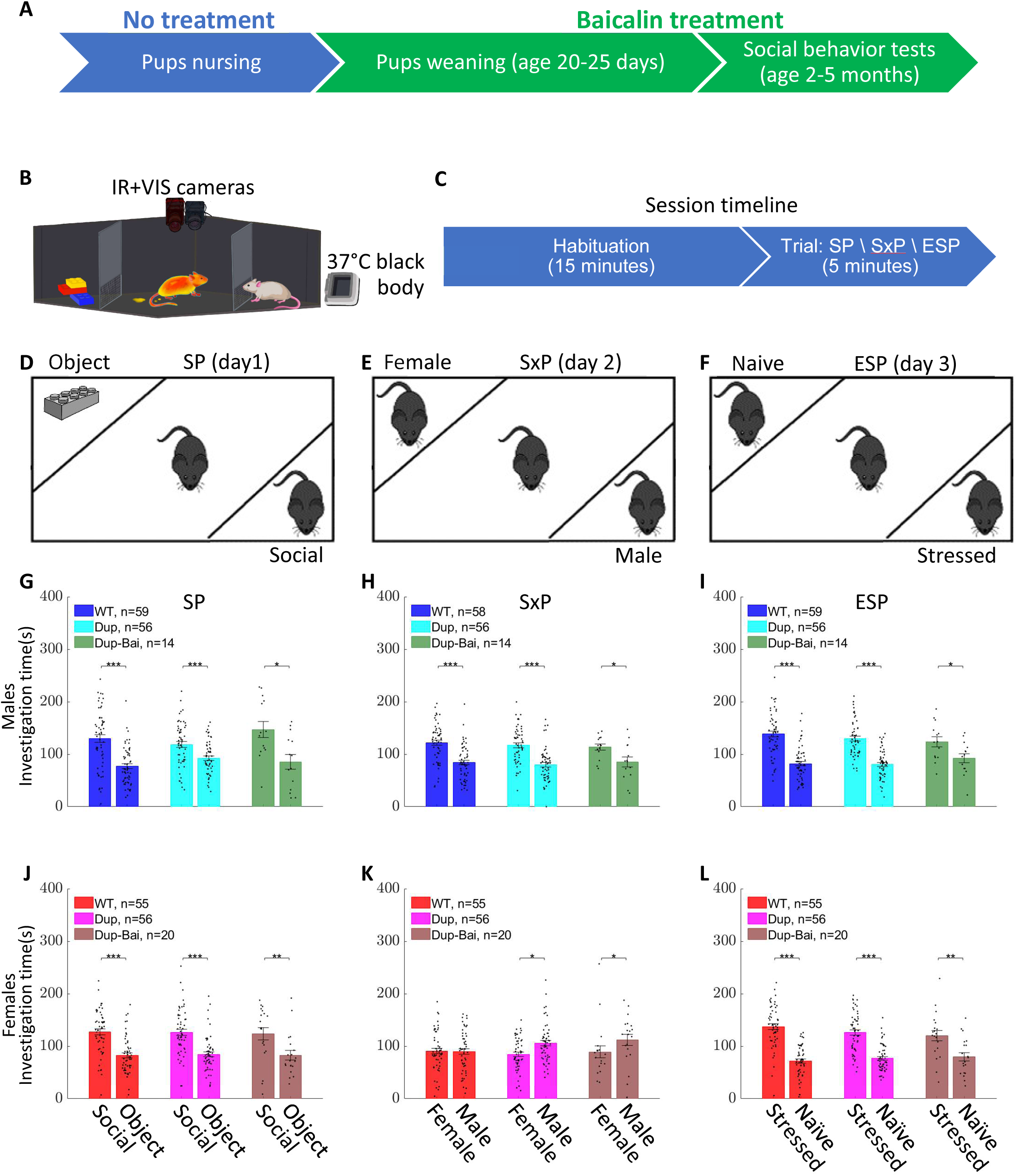
Experiment timeline, setup, and mice’s social discrimination behavior. Experiment timeline (**A**) includes treating the Dup and WT mice with Baicalin dissolved in their drinking water (0.15mg/ml) since weaning. Experiment setup (**B**) includes an arena imaged by both a visible light camera and a thermal camera. Each test timeline (**C**) includes 15 minutes of habituation followed by 5 minutes of a trial. In the SP trial, which is performed on day 1 (**D**), animals were exposed to a sex-matched mouse and a Lego toy. In the SxP trial, which is done on day 2 (**E**), animals were exposed to a female mouse and a male mouse. In the ESP test, which is done on day 3 (**F**), animals were exposed to a stressed mouse and a naïve mouse. (**G-I**) show the mean time (±SEM) spent by the three male groups on investigating stimulus1 (left bar in each pair) and stimulus2 (right bar in each pair) during SP (**G**), SxP (**H**), and ESP (**I**). Stimulus1 is defined as the social stimulus in SP, the female in SxP, and the stressed mouse in ESP. (**J-L**) Shows the same as (**G-I**), for female mice. *p<0.05, **p<0.01, ***p<0.001, using FDR corrected (3 comparisons per panel) two-sided Wilcoxon rank sum test.

### Setup and Video Acquisition

Our setup is based on the setup used in [29] and [30]. Briefly, a black plexiglass box arena (37cm x 22 cm x 35 cm) was placed in a sound-attenuated chamber. An infrared (IR) thermal camera (Opgal’s Thermapp MD with 8.66 frames per second) and a visible light camera (either Flea3 or Grasshopper3, both by Teledyne FLIR) with a wide-angle lens and acquisition rate of 30 frames per second, were placed above the arena (about 70 cm above the arena’s floor). The IR camera is designed to measure human skin temperature and outputs apparent temperature per pixel. Video was captured using the manufacturer’s SDK (SDK version: EyeR-op-SDK-x86-64-2.15.915.8688-MD) and python scripts previously published in [30]. To improve image quality and pixel uniformity, the IR camera was turned on at least 15 minutes before starting the experiment. A set of images of the uniform surface were captured before each experiment and were later used for pixel non-uniformity correction using the following protocol: 16 images of the uniform surface were captured and averaged, the mean of the average image was subtracted to get a zero-mean non-uniformity image, which was later subtracted from each image in the recorded IR video. A high-emissivity black body (Nightingale BTR-03, by Santa Barbara Infrared Inc.) was set to 37°C and was placed in the field of view of the IR camera. The black body image was used to compute a correction offset for the apparent temperature measured by the camera. The experimental setup is illustrated in Fig. 1B.

### Social behavior tests

We used three social discrimination tests, each performed on a different day and with a similar protocol to the one in [31]. All experiments included a 15-minute habituation period in which the mouse got used to the arena, followed by a 5-minute trial period in which two different stimuli were introduced to the arena (Fig. 1C). During the habituation period, two empty triangular chambers were located in two randomly chosen opposite corners of the arena. Each chamber has a metal mesh (18cm width x 6cm height with 1×1 cm holes) at its bottom, to allow the subject mouse to investigate the chamber’s content. After the end of habituation, the empty chambers were replaced with stimulus-containing chambers, and the trial started. During the Social Preference test (SP), one chamber contained a sex-matched mouse while the other contained a Lego toy. During the Sex Preference test (SxP), one chamber contained a male mouse while the other contained a female mouse. During the Emotional State Preference test (ESP), one chamber contained a sex-matched stressed mouse (restrained in a 50-ml plastic tube for 15 min before the test), and the second chamber contained a sex-matched naïve mouse (Fig. 1D-F). All social stimuli were age-matched and novel to the subject mouse.

### Analysis of investigation time

Analysis of the time dedicated by the subject mouse for investigating each stimulus (investigation time) was done using the videos recorded by the visible light (VIS) camera (Fig. 1B). These videos were analyzed using the TrackRodnet software as previously described in [29]. The software computed the investigation time for each stimulus, as well as the relative discrimination index (RDI), defined as the difference in investigation time between the two stimuli, divided by their sum. Positive RDI reflects a preference towards stimulus 1, defined as the social stimulus in the SP test, the female stimulus in the SxP test, and the stressed mouse stimulus in the ESP test. Negative RDI reflects a preference towards the object stimulus in the SP test, the male stimulus in the SxP test, and the naïve stimulus in the ESP test.

### Annotating the thermal videos

We followed the same procedure as previously described in [30]. A previously described GUI [30] was used to annotate the arena’s floor, mark the side of each stimulus, mark the location of the black body surface, and specify the first and last frames of the habituation period and the trial period. In addition, a location on the plastic shelf that holds the arena was marked as an ambient temperature reference point and was used to compensate for the room temperature (Fig. 2A).

**Figure 2:**
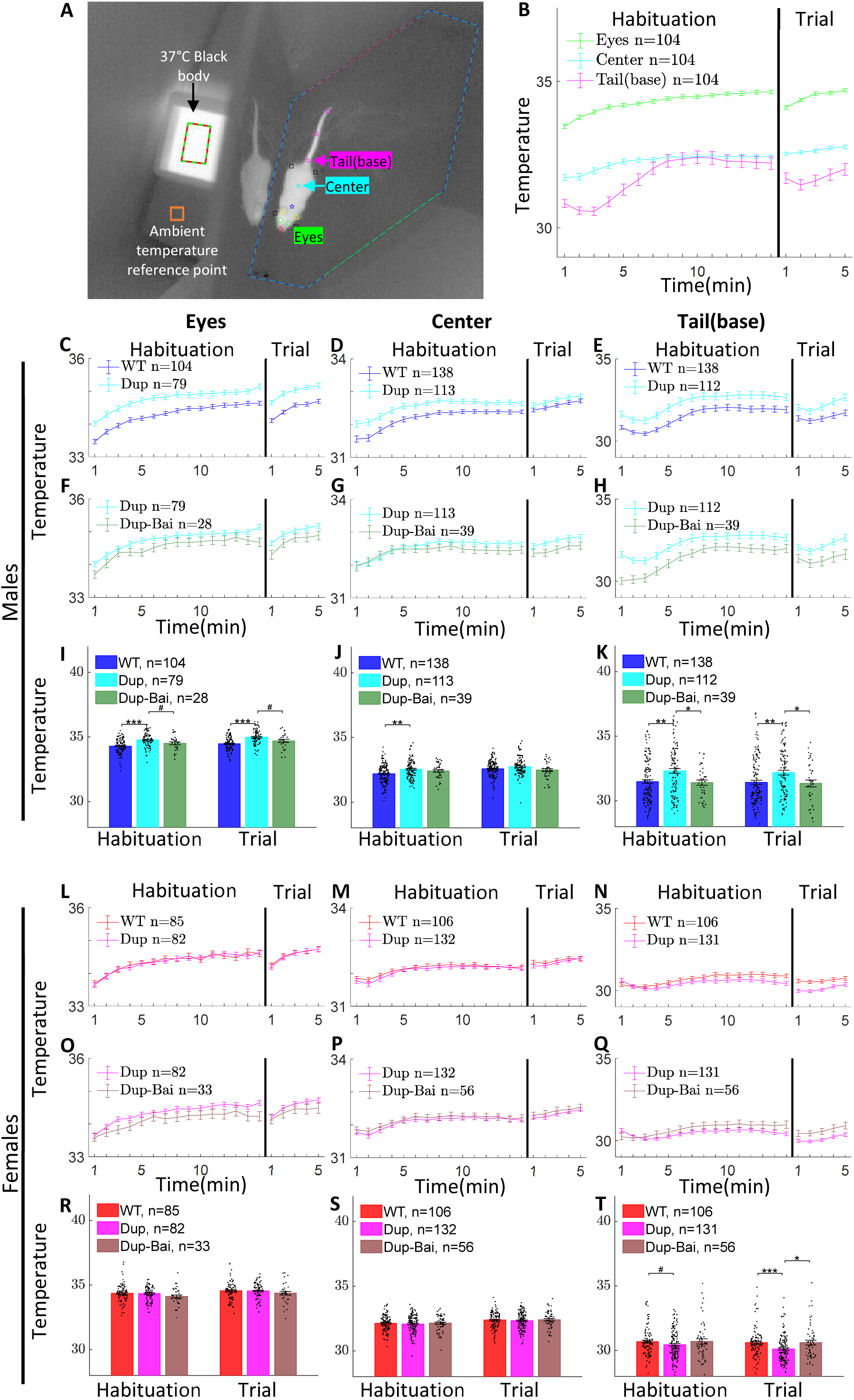
Baicalin treatment ameliorates the surface temperature phenotype of Dup mice. Thermal image (**A**) overlayed with the manually marked arena’s floor (blue-cyan line), stimulus sides (green line for stimulus1 and red line for stimulus2), blackbody 37°C surface (green-red polygon), ambient temperature reference point (orange rectangle), and automatically detected key points using the DLC algorithm. These key points include the Eyes, Center, Tail(base), as well as the nose, four limbs (black rectangles), tail middle and tail end (pink stars), and ears (yellow triangles). (B) shows the mean (±SEM) surface temperature dynamics of the Eyes, Center, and Tail(base) for WT male mice (SP, SxP and ESP sessions were pooled together). Comparisons of the surface temperature dynamics in the Eyes, Center, and Tail(base) are shown separately for WT vs Dup and Dup vs Dup-Bai in (**C-E**) and (**F-H**), respectively, and are quantitatively analyzed in (**I-K**). Bars show the mean temperature ±SEM. (**C-K**) include data from SP, SxP, and ESP tests pooled together. (**L-T**) shows the same for female mice. Comparisons of WT to Dup and Dup to Dup-Bai in (**I-K**) and (**R-T**) were done using the two-sided Wilcoxon rank sum test and were FDR corrected (4 comparisons per panel).

**Figure 3:**
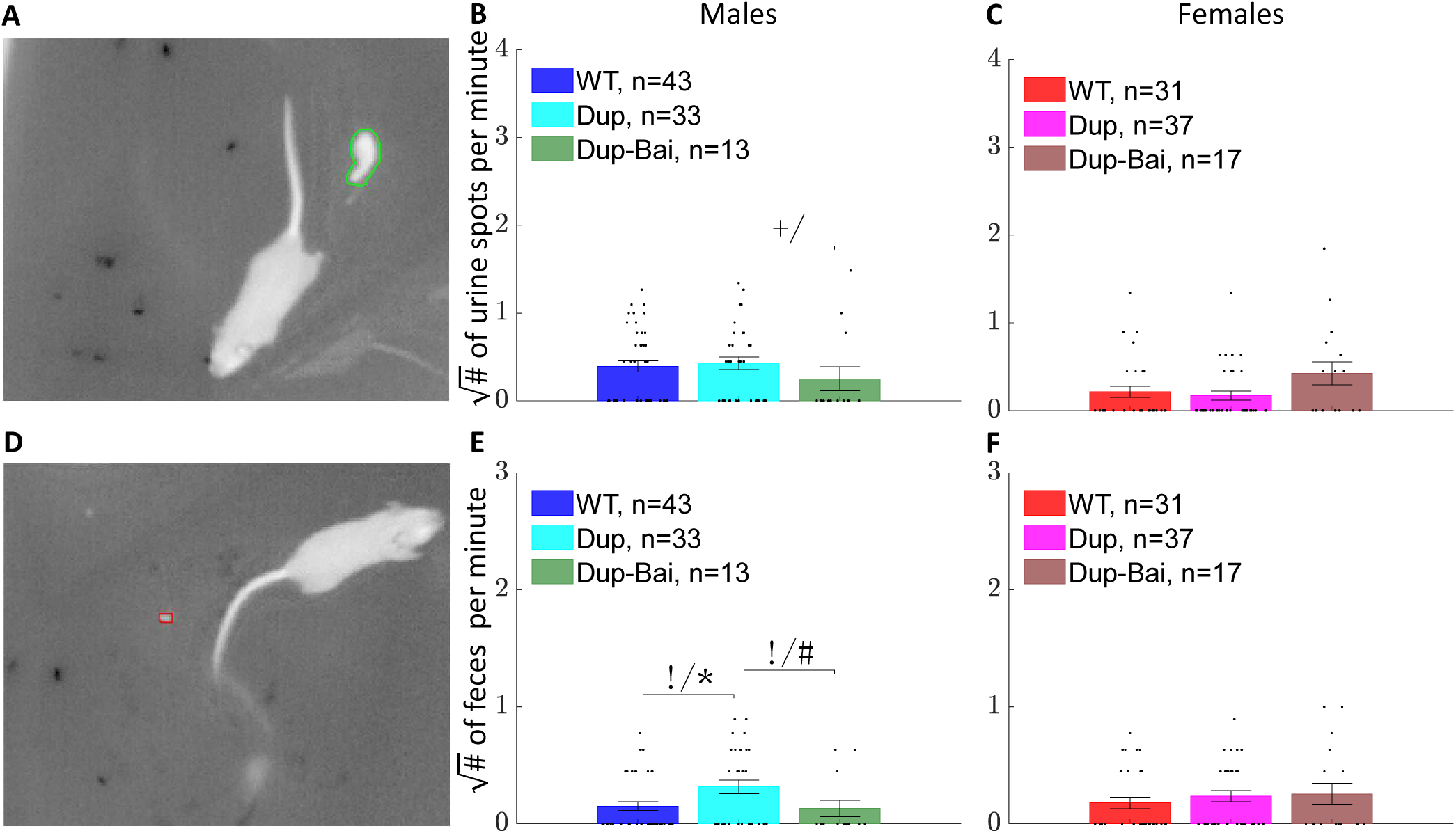
Baicalin ameliorates the higher defecation rate in Dup male mice during the trial period of the SP test. Automatic detection of a urine spot (**A**) and a fecal deposition (**D**) by the DeePosit algorithm (see Methods) in a thermal video of a Dup male mouse during the SP test. The urine and fecal deposit detections are marked with a green line and a red line, respectively. The mean (Square root) number of urine spots per minute ±SEM in males (**B**), and Females (**C**), and of fecal deposits per minute ±SEM in males (**E**) and females (**F**) were compared between WT and Dup, and between Dup and Dup-Bai groups. FDR-corrected (2 comparisons per panel) pairwise comparisons using two-sided Wilcoxon ranksum test were marked with #,* for p<0.1 and p<0.05, respectively. Pairwise comparisons using a two-way Chi-square test were used to compare the distribution of zeros between each pair of compared groups. Significance in the Chi-square test (FDR-corrected, 2 comparisons per panel) was marked with !,+ for p<0.1 and p<0.05, respectively.

### Analysis of micturition and defecation activities

Micturition and defecation rates (spots per minute) were computed using DeePosit software, as previously described [30]. Some unreliable IR videos were excluded (64 of 743 videos were excluded, see Supplementary Table 1), mostly due to non-optimal positioning of the arena below the camera, which resulted in the occlusion of part of the arena floor. We considered an occlusion to be significant (and hence, excluded the video) if part of the mouse’s body (and not just its head) is occluded when the mouse faces one of the stimuli.

### Analysis of body surface temperature

Analysis of the surface temperature of the subject mice was done in the same manner as described in [8]. Briefly, key points marking the subject mouse body parts were automatically detected in the thermal videos using DeepLabCut (DLC) software [32] version 3.0.0rc4, which was trained on 1999 manually marked IR images. The key points included the nose, left and right eye, left and right ear, neck, center (the body center), the four limbs, and three points on the tail: tail-base, tail-middle and tail-end (see Fig. 2A). To avoid DLC from identifying key points on the stimulus mouse’s body (which was visible during certain trials), the region containing the stimulus (defined as the region that is 14 pixels away from the lines that mark the stimulus chamber’s border) was replaced with a constant value equivalent to 22°C.

To ensure the reliability of the tracked key points, only frames in which the DLC grade for the key points: neck, center, and tail-base were all higher than 0.7 (grades are between 0 and 1) were considered for further analysis. Furthermore, a key point was considered valid in a specific frame only if its DLC grade is higher than 0.8 and it is located inside the borders of the annotated arena floor and at least 6 pixels away from its borders (to account for the spatial filters detailed below). In addition, the eyes and nose key points were considered valid only if all three of them were valid and with DLC grade > 0.8. Tail-base temperature was defined as the maximal temperature within a radius of 6 pixels to compensate for possible minor mistakes in the detected location. Similarly, for the eyes, the maximal temperature in a radius of 3 pixels was used, and the maximal temperature for the left and right eyes was used as the eyes’ temperature in the analysis. In order to reduce noise, we used a smoothed image (Gaussian filter of 11×11 pixels and std=2.5 pixels) for extracting the temperature of the body center. We corrected the measured temperature to compensate for the ambient temperature in the room, as detailed in [8]. Finally, the median temperature over each minute of the habituation and trial was computed. For periods longer than one minute, the average of the medians of the relevant minutes was used. Videos in which the mouse had a dark spot on its fur (due to urine or feces on its fur), or fur damage were excluded (32 out of 743 videos were excluded, see Supplementary Table 1).

### Animal experiment dates and batches

To increase the statistical power, we unified data from three batches of the mice experiments. Batch1 and Batch2 include recordings from WT and Dup mice, but do not include a Baicalin treatment group. Batch3 includes recordings from additional non-treated mice and from Baicalin-treated mice. Batch1 was recorded between April 2023 and March 2024. Batch2 was recorded between December 2024 and January 2025, and Batch3 was recorded between March 2025 and August 2025.

### Patient-derived neurons

We used the cell lines UOHi010 (Control) and UOHi011 (7Dup), which were previously described in [33]. Briefly, patient-derived neurons were generated from blood samples of a 7.5-year-old male child (UOHi011) and his father (age 39) as a control (UOHi010). The child was diagnosed with ASD, obsessive compulsive disorder (OCD), social anxiety, selective mutism, and a ∼1.8Mb duplication in 7q11.23 section that includes the GTF2I gene: arr 7q11.23(72,722,981-74,489,889)x3 GRCh37, diagnosed by Illumina SNP array GxG_ComprehensiveArray_V1. Experiments were approved by the University of Haifa Ethics Committee for Experiments in Humans (approval No. 526/20). See [33] for the full details of the reprogramming protocol. Briefly, a regular blood test from the patient and his father was collected into heparin-coated tubes. Peripheral blood mononuclear cells (PBMCs) were isolated from the blood samples and stored in 10% fetal bovine serum (FBS, F7524 Merck) DMSO solution, and kept in liquid nitrogen. CytoTune-iPS Sendai Reprogramming Kit (A16518, Life Technologies, Carlsbad, CA, USA) was used for reprogramming the isolated PBMCs into iPSCs. Embryonic bodies (EBs) were generated from iPSCs and plated onto poly-l-ornithine/laminin-coated plates and fed with complete Neural Progenitor Cell (NPC) medium (DMEM/F12 with Glutamax (1:100), B27 supplement with RA (1:100), N2 supplement (1:100), Laminin (1 mg/ml), and 20 ng/ml FGF2) until reaching full confluency. NPCs were replated one day later and differentiated into cortical pan-neurons by feeding with differentiation medium containing DMEM/F12, N2, B27, Glutamax, ascorbic acid (200 nM), cyclic AMP (cAMP; 500 mg/ml), laminin (1 mg/ml), BDNF (20 ng/ml), and GDNF (20 ng/ml) for 10 days. Between days 11 and 14, the cells were dissociated again and then fed with Brainphys medium with B27 supplement (1:100), N2 supplement (1:100), ascorbic acid (200 nM), cAMP (500 mg/ml), BDNF (20 ng/ml), GDNF (20 ng/ml), and laminin (1 mg/ml).

### Treating cells with Baicalin

Baicalin was dissolved in DMSO (10mg/ml) and filtered to remove possible pathogens. 0.5ul/ml of the Baicalin dissolved in DMSO was added to the treated cells (7Dup-Bai group) feeding media starting from day 16 of the differentiation from NPCs to neurons, resulting in 5ug/ml (11.2 uM) Baicalin concentration in the cells’ media. Control cell line, as well as the non-treated patient cell line (7Dup line), received just the DMSO (0.5ul/ml) without Baicalin from day 16 until the end of the experiment.

### Whole-cell patch clamp recordings

Whole-cell patch clamp recordings were done in the same protocol as in [33], and at two time points (TPs): at days 33-38 (TP1) and days 74-79 (TP2) after the start of differentiation from NPC to neurons. Culture coverslips were extracted from the 48-well plate and put on a microscope stage within HEPES-based artificial cerebrospinal fluid (ACSF) pre-warmed to 37°C containing (in mM): 10 HEPES, 139 NaCl, 4 KCl, 2 CaCl2, 10 D-glucose, 1MgCl2 (pH 7.4, osmolarity adjusted to 310 mOsm), and Baicalin (dissolved in DMSO and in the same concentration as in feeding media) for the 7Dup-Bai group or just the DMSO (0.5ul/ml) for the Control and 7Dup groups. The recording micropipettes were filled with an internal solution containing (in mM): 130 K-gluconate, 6 KCl, 4 NaCl, 10 Na-HEPES, 0.2 K-EGTA, 0.3 GTP, 2 Mg-ATP, 0.2 cAMP, 10 D-glucose (pH 7.4, osmolarity adjusted to 290–300 mOsm). The recording was done at room temperature using Clampex v11.1 with a sampling rate of 20 kHz. The experiment duration for each coverslip was limited to 3 hours. Four coverslips from each of the three groups (Control, 7Dup, 7Dup-Bai) were used for patch clamp at each time point (24 coverslips in total). Five recordings were done sequentially per cell, at the first and last (fifth) recording, cell voltage was clamped to −60 mV, and the currents injected into the cell in order to maintain the voltage clamp were recorded. Cells for which the absolute mean current surpassed 70 pA were ignored. Recording length was 1 minute. These recordings were used for the analysis of excitatory postsynaptic currents (EPSCs). In the second recording (which is not analyzed in this work), voltage clamp steps of 400ms from −100 to 90 mV were carried out, and the currents were recorded. In the third recording, neurons were held in current clamp mode. Current was adjusted to reach a cell voltage of approximately −45 mV. A 56-second recording of the cell voltage was carried out and was later used to compute the number of spontaneous action potentials. In the fourth recording, the neurons were held with a constant current manually adjusted (base current) to bring the cell to −60 mV. Next, a series of 38 current injections with steps of 6 pA starting from −10 pA below the base current up to 218 pA were injected for periods of 400ms. This recording was used to quantify the number of evoked action potentials (only the first 10 current steps were analyzed).

### Automatic detection of excitatory postsynaptic currents (EPSCs)

Automatic detection of EPSCs (Fig. 4J and Supplementary Fig. 6) was done using a template matching algorithm, which was implemented in Matlab and consists of the following steps. Code will be published at https://github.com/davidpl2/BaicalinFor7Dup.

**Figure 4:**
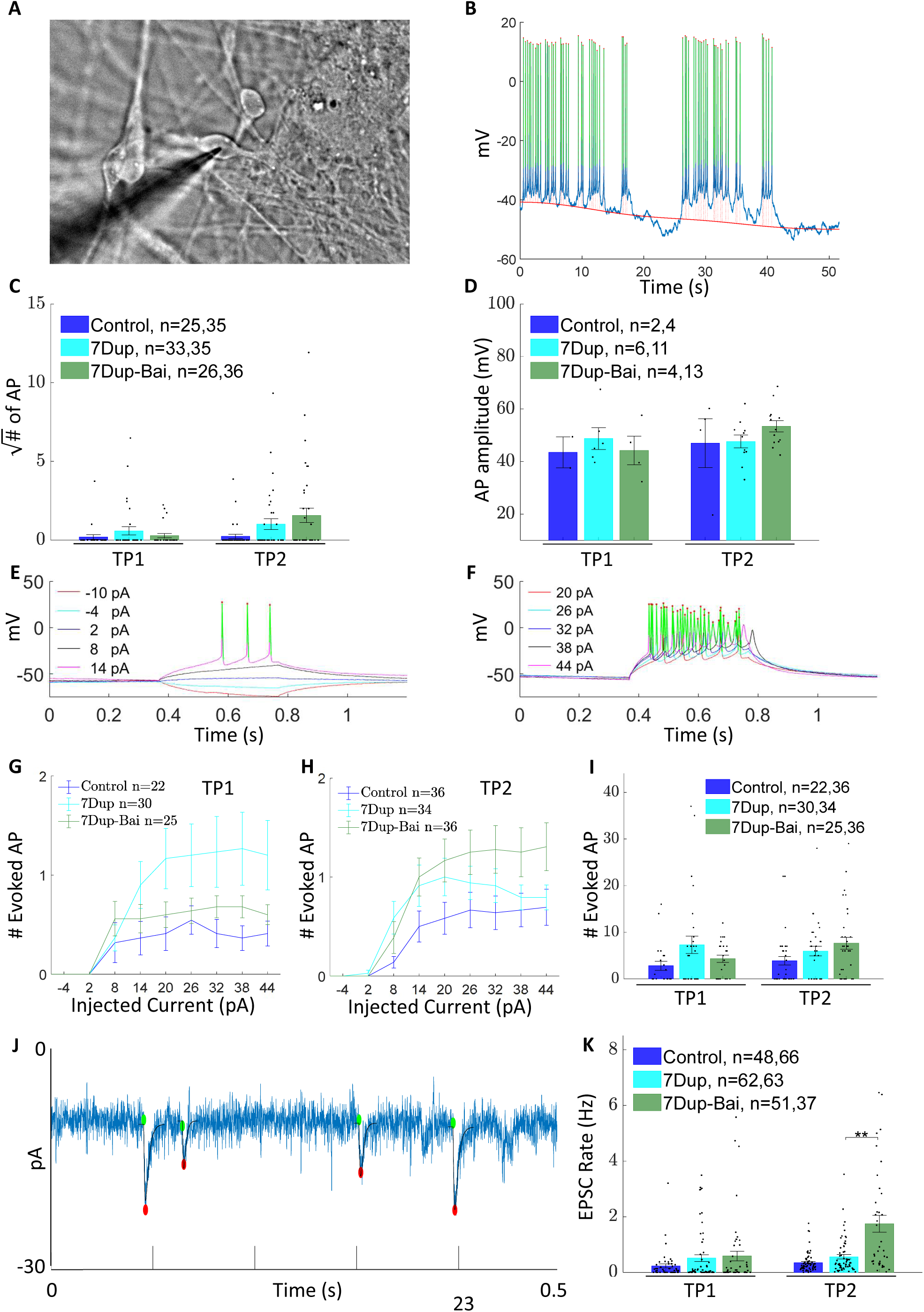
Baicalin increases EPSC rate in 7Dup patient-derived neurons. (**A**) Brightfield image of a patched cell (from 7Dup-Bai cell culture at time point 2). (**B**) Shows an illustration of automatic detection of spontaneous action potential in current clamp mode. Before recording, the current was adjusted to bring the cell to an initial voltage of approximately −45mV. Detected action potentials are marked in green, and the maximum of each action potential is marked with a red dot. Baseline computation (red line) was used for the computation of the AP amplitudes, which are marked with dashed red lines. No significant difference was found in the # of AP or in the amplitude of the Baicalin-treated cells (**C,D**). (**E,F**): Example of automatic detection of evoked action potentials. In each step, a different current was injected into the cell. (**E**) shows the AP detection in the first five current steps (−10 to 14 pA). (**F**) shows steps 6-10 (20 pA to 44 pA). Detected action potentials are marked in green, and red points mark the peaks. (**G,H**) shows the mean number of evoked AP ±SEM as a function of the injected current for the Control, 7Dup, and 7Dup-Bai cells in time point 1 (**G**) and time point 2 (**H**). Though some trends are visible, no statistically significant difference was found (**I**). (**J**) Example of automatic EPSCs detection in a patch clamp recording using voltage clamp. Each detected EPSC is marked with a green dot at its onset, a red dot at its peak, and a black line that shows the matching synthetic EPSC profile that was found using template matching. See Methods for details. The panel shows a 0.5-second period of a 60-second recording. In the 2^nd^ time point (TP2), Baicalin-treated patient-derived neurons show elevated EPSC rate (**K**). Each point in (**K**) represents a single recording (each cell was recorded twice, see Methods). Panels (**C,D,I,K**) show mean value ±SEM. Comparisons between WT and 7Dup and between 7Dup and 7Dup-Bai were done using two sided Wilcoxon rank-sum test and were FDR corrected (4 comparisons per panel).

#### Synthetic EPSCs profile generation

Synthetic EPSCs profiles (shown in Supplementary Fig. 6A) were generated by summing two exponential functions, denoted *expRise* and *expDecay*.

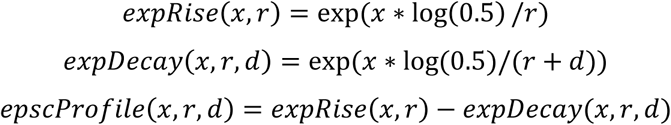

where x is the index of the time (starts at 0), r and d are parameters that together determine the rise and decay time of the generated EPSC profile, and log is the natural logarithm function. Intuitively, both *expRise* and *expDecay* are equal to 1 at x=0, hence *epscProfile*(0,r,d)=0. However, *expRise* decays faster and reaches the value of 0.5 at x=r, while *expDecay* decays more slowly and reaches the value 0.5 at x=r+d. We generated 168 synthetic EPSC profiles by choosing 12 values for r and 14 values for d, and then using all combinations of r and d values. The smallest r value was selected as 0.15 milliseconds and was iteratively increased by 30% each time to create the additional 11 values. Similarly, the smallest d value was selected to be 0.5 milliseconds and was iteratively increased by 30% each time to create the other 13 values. Finally, the profiles were padded with 5ms of a constant zero value at their beginning. The rise time and decay time of the generated EPSCs profiles are shown in Supplementary Fig. 6B. The rise time is defined as the time from the onset of the EPSC until reaching the peak’s minimal value. The decay time is defined as the time from the peak until rising back to 2/3 of the decline (Supplementary Fig. 6B). The profile length was set to include almost all of the decay period (until reaching a value higher than −0.02) but up to a maximal length of 1201 samples (60 milliseconds).

#### Preprocessing

As some of the recordings had noise in 50Hz and 60Hz frequencies (probably due to the power outlet AC frequency), these frequencies were removed by transforming the recorded signal to the Fourier domain, zeroing the Fourier coefficients that match 50Hz, 60Hz, and their 100Hz and 120Hz harmonics, and transforming back to the time domain using inverse Fourier transform. The resulting signal went through additional smoothing with an 11-element (0.55 millisecond) Gaussian kernel with a sigma of 1.325 elements (0.066 millisecond). Note that EPSC detected in the first 0.5 second and last 0.5 second of the recording were ignored since the preprocessing result is less accurate in these periods.

#### Template Matching

A normalized cross-correlation (Pearson correlation) was computed between each of the 168 synthetic profiles and each position in the recorded signal. The maximal correlation value for each position in the signal was saved. Positions in the recorded signal for which: a. their correlation score is higher than 0.65 and are local maxima in the range of [−5..+5] milliseconds were defined as detection candidates. The rise amplitude of each detection candidate was defined as the difference of the median value during the 5 milliseconds before the current drop (drop location is defined by the matched synthetic profile, i.e. the synthetic profile with the highest correlation) and the peak value (defined as the minimal value of the signal in a range of ±1millisecond from the peak location according to the matched synthetic profile). The decay amplitude of a detection candidate was defined as the maximal signal after the peak (2.5ms .. 32.5ms after the peak) minus the peak value. Detection candidates matching the following conditions were considered as EPSC detections: a. a rise amplitude greater than 5 pA, and greater than 0.3 of the decay amplitude, and greater than 2\**local_std* (computation is described next), b. decay amplitude higher than 0.5 of the rise amplitude. The *local_mean* and *local_std* were computed using a sliding window of 1-second width. After the first computation, the local mean was subtracted from the signal, and values that exceeded ±3\**local_std* were clamped to ±3\**local_std*, respectively. The *local_std* was re-computed on the clamped signal. This clamping mitigates the effect of outliers and EPSCs on the computed *local_std*. Examples of detected EPSC and their matched synthetic profiles are shown in (Supplementary Fig. 6C,D, and Fig. 4J). Detections of EPSC that had a rise amplitude higher than 30 pA were ignored in the analysis, as they might be a result of an action potential and not an EPSC.

### Automatic detection of action potentials (AP) in spontaneous AP recording and Evoked AP recording

We used a custom peak detector for detecting action potentials (AP) in the recordings of spontaneous AP and evoked AP. Code will be published (https://github.com/davidpl2/BaicalinFor7Dup). The peak detection algorithm checks all possible start and end points in the recorded signal and determines if the interval between the start and end points is a valid action potential. Action potentials are then sorted by their grade (denoted *AP_Grade_*) and added to the final list of detected action potentials as long as they don’t overlap with another action potential that is already in that list. Valid action potentials were defined as intervals for which the following preliminary conditions are met:

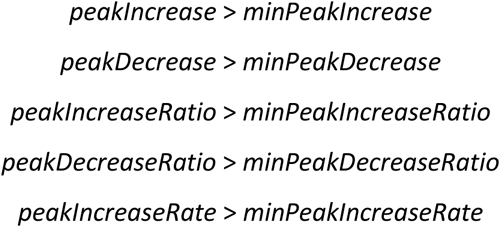

Where on the right side are the algorithm’s parameters, and where:

*leftBorderVal, rightBorderVal* = the value at the start and end point of the interval

*maxVal* = the highest value in the current interval

*peakIncrease* = *maxVal* - *leftBorderVal*;

*peakDecrease* = *maxVal* - *rightBorderVal*;

*dRange* = the dynamic range of the signal, defined as its maximum minus its minimum.

*peakIncreaseRatio* = *peakIncrease* / max(1e-6,*dRange*);

*peakDecreaseRatio* = *peakDecrease* / max(1e-6,*peakIncrease*);

*maxValInd* = index of the maximal value in the interval (ranging from 1 to the interval length)

*peakIncreaseRate* = *peakIncrease* / *maxValInd*;

Intervals that met the preliminary conditions received an *AP_Grade_* that was used to prevent overlapping of detected AP (as previously described). The grade was designed to give priority to sharp peaks:

*AP_Grade_* = *peakIncreaseRatio*^2^ ∗ *peakDecreaseRatio*^2^ ∗ *peakIncreaseRate* ∗ *peakDecreaseRate*

Where *peakIncreaseRatio*, *peakIncreaseRate*, and *peakDecreaseRate* were defined above, and:

*intervalLength* = the length of the current interval

*peakDecreaseRate* = *peakDecrease* / (*intervalLength*-*maxValInd*+1);

The parameters that were used for the automatic detection of AP were:

*minPeakIncrease* = *minPeakDecrease* = 10; (units = mV)

*minPeakIncreaseRatio* = 0.1; (units = % of the dynamic range)

*minPeakIncreaseRate* = *minPeakIncrease*/100; (units = mV per 1/20000 sec)

*minPeakWidth* = 100; (= 5ms)

*maxPeakWidth* = 500; (= 25ms)

*minPeakDecreaseRatio* = 0.6; (units = fraction of the increase)

#### Baseline computation

baseline computation was used for computing the amplitude of the AP in the spontaneous AP recording. AP amplitude was defined as the peak value minus the baseline value (See Fig. 4B). The baseline of a signal *y* was computed by:

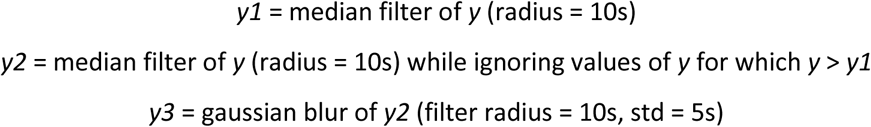

### Calcium Imaging

Calcium imaging of the three groups of cells (Control, 7Dup, 7Dup-Bai) was done at two time points (TPs): TP1 (days 33-37) and TP2 (days 74-79) to assess the neuronal activity of the cell cultures by monitoring the cells’ calcium activity. Cells’ coverslips were incubated in a 37°C incubator for one hour in 2 mM Fluo-5 AM fluorescent calcium indicator, which was dissolved in HEPES-based ACSF, containing (in mM): 10 HEPES,139 NaCl, 4 KCl, 2 CaCl2, 10 D-glucose, 1MgCl2 (pH 7.4, osmolarity adjusted to 310 mOsm), and Baicalin (dissolved in DMSO and in the same concentration as in feeding media) for the 7Dup-Bai group or just the DMSO (at the same concentration as in feeding media) for the Control and 7Dup groups. After the incubation, the cover slips were transferred to a fresh ACSF with the same composition except for the Fluo-5 AM, which was excluded. ACSF was pre-warmed to 37°C. Microscopy videos were recorded using a Leica Thunder imager microscope and LAS X version 3.7.5.24914 software. During recording, cells were kept at 37°C using an incubator. Video recording was done for ∼10 regions in each coverslip, at x200 magnification, 2048×2048 pixels per grayscale image, capturing a 655.23×655.23 micrometer region (0.32 micrometers per pixel), with a frame rate of 10 frames per second and for 12 minutes per region (only the first 10 minutes were analyzed). Non-compressed raw images were recorded and analyzed using custom Matlab scripts.

### Automatic analysis of calcium imaging recordings

Calcium imaging videos were analyzed by custom Matlab-implemented heuristic algorithms. The code will be published (https://github.com/davidpl2/BaicalinFor7Dup). The heuristic algorithm is comprised of the following steps.

#### Automatic detection of active cell

The active cell segmentation algorithm automatically clusters together pixels that are active (show calcium transients), correlated (and hence could belong to the same cell), and that form a spatially connected region. In detail, videos were smoothed to reduce noise (box filter with spatial radius of 6 pixels and temporal radius of 2 images), which generated a smooth calcium trace (*SCT*) for each pixel. Dark level was set as the minimal intensity in the smoothed video minus 1 gray level (raw gray level range is 0. .2^16^ − 1 gray levels). A high pass *HP(p,t)* filter was computed for each pixel p at time t by subtracting *LP(p,t)* which is defined as the mean intensity of pixel p during [t-10s .. t+10s]. Denote *NormHP(p,t)* as the normalized version of *HP(p,t)* computed for each pixel p by:

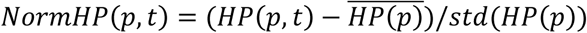

Deonte the high pass to low pass ratio *HtoL*:

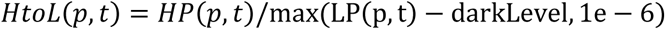

The maximal *HtoL* per pixel p over all frames was computed (denoted *maxHtoL(p)*, illustrated in Fig. 5A). A pixel p was considered active if *maxHtoL(p)* was 3 times higher than the 3% percentile of the *maxHtoL* (this percentile is an estimation of the noise level in non-active pixels in *maxHtoL*). A mask of active pixels (denoted *fgMask*) was initialized by setting it to 1 for active pixels and 0 elsewhere. An iterative algorithm extracted active pixel clusters from *fgMask* by the following steps. Let p be the pixel with the maximal *maxHtoL* in the eroded (radius=3 pixels) version of *fgMask* (erosion was done to ignore very small clusters). Let *t* be the time at which *p* achieves its maximal *HP* value. Let *blobMask* be a mask image that equals 1 for pixels which have *NormHP* value larger than *0.6*NormHP(p,t)* at time *t* and that are included in *fgMask* (i.e., pixels with a significant calcium transient in comparison to pixel *p* in time *t* and that are still relevant for clustering). The normalized cross-correlation between the *NormHP* values of *p* and the *NormHP* values of all other pixels in *blobMask* is computed. Pixels with correlation smaller than 0.9 are removed from *blobMask*, as these pixels probably do not belong to the same cell as *p*. If *blobMask* has fewer than 500 pixels (a very small cluster or several very small clusters), then this iteration will not generate a new cell segment and the pixels in the dilated (radius=4 pixels) version of *blobMask* are removed from *fgMask*. Otherwise, *blobMask* goes through a morphological close operation (radius=2) that closes small holes in the mask, and a connected region that includes *p* is extracted from it. The pixels in this cluster (and their close neighbors) will not be considered for clustering in the next iterations by setting *fgMask* to 0 for the pixels in the dilated version of this cluster (radius = 4). To ensure that there is no overlap between cells, pixels that were previously associated with active cells are removed from the currently detected cluster. If these cause the cluster to split, then the part of the cluster that contains pixel *p* will be used in the next steps. If the cell cluster is larger than 500 pixels and its minimal width (in any direction) is at least 21 pixels (ignoring thin elements that usually belong to neurites), then a new cell cluster is added to the list of cell clusters, and *p* is considered the representative pixel of this cluster. This process continues as long as there are non-zeroes in *fgMask*.

**Figure 5:**
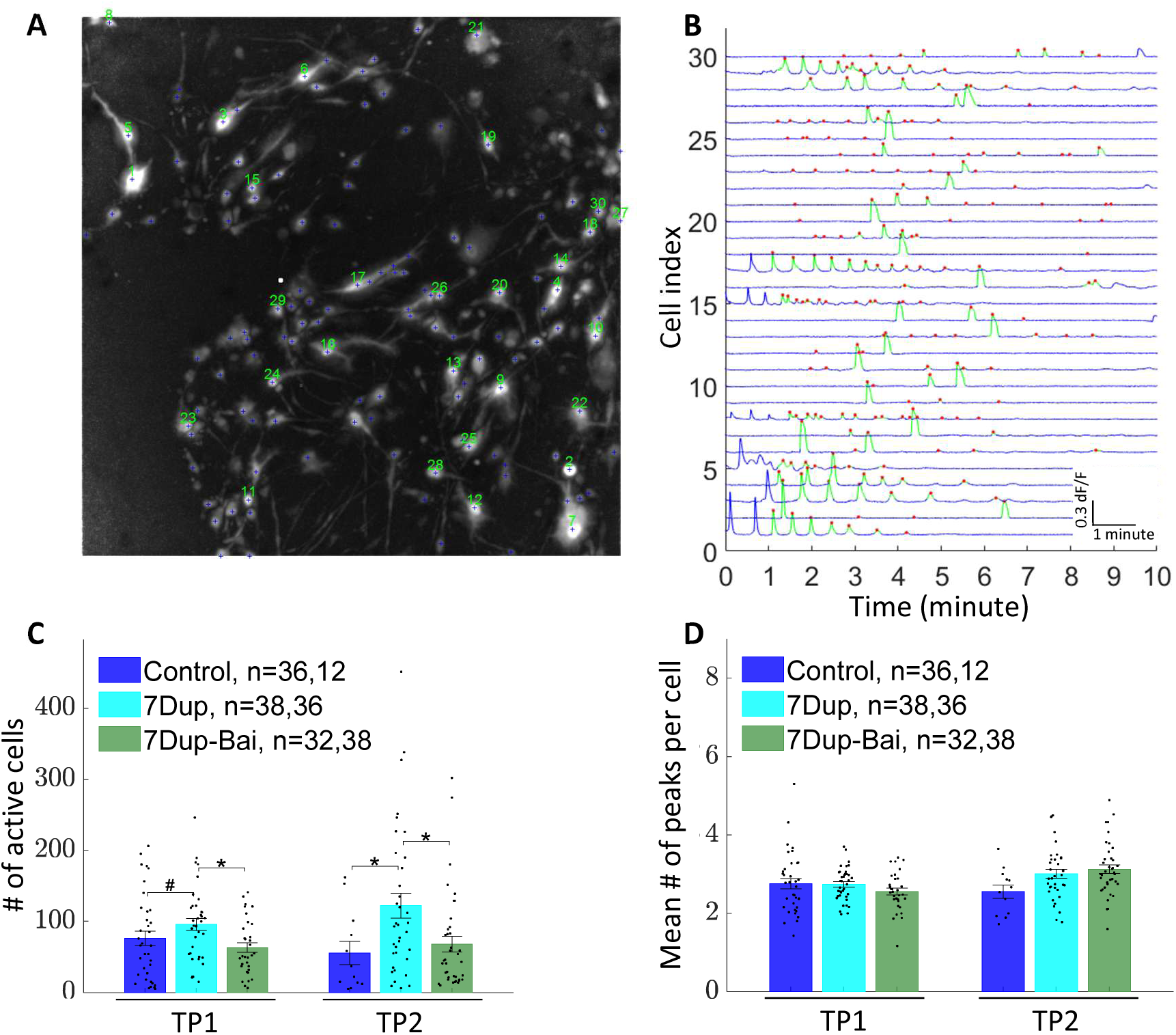
Baicalin reduces the number of active cells in 7Dup patient-derived neuron culture. (**A**) Example of calcium activity image (maxHtoL, see Methods) overlayed by the automatically detected active cells marked with a blue + sign. Processed fluorescence traces (dF/F) of active cells 1-30 (marked with green text in (**A**)) are shown in (**B**). Each automatically detected calcium transient in (**B**) is marked with a green line and a red dot that marks its peak. No detections were made in the first and last minute of the recordings (see Methods for details). (**C**) Mean number ±SEM of active cells in calcium video recording during the most active 2 minutes of each recording. Baicalin-treated cells showed a lower number of active units per region during time points 1 and 2. The mean number ±SEM of calcium transients per cell during the most active 2 minutes of each recording is shown in (**D**). Comparisons between WT and 7Dup and between 7Dup and 7Dup-Bai in (**C,D**) were done using a two-sided Wilcoxon rank-sum test and were FDR corrected (4 comparisons per panel).

#### Automatic peak detection in calcium transients of representative pixels

Peak detection relies on the smooth calcium traces (*SCT*) defined in the previous section for the representative pixels. We define dF/F values per representative pixel *p* at time *t* by computing:

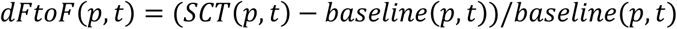

The baseline computation comprises the following steps: a. A median filter of *SCT* is computed across time (radius=35s). b. A median filter is computed (radius = 35s) on SCT again, but this time, ignoring values of SCT that surpass their first median filter result, hence, minimizing the effect of the calcium transient peaks on the computed baseline. Next, the result is further smoothed by a Gaussian filter (radius of 70s and a sigma of 35s) to get the baseline trace.

To find calcium transients, a threshold is computed for each representative pixel p in *dFtoF* by the following method: first, the std of *dFtoF(p)* across time is computed (denoted *std_1_(p)*). A second std computation (denoted *std_2_(p)*) of *dFtoF(p)* across time is computed while ignoring values in *dFtoF(p)* with absolute value larger than 2\**std_1_(p)*. The std of *dFtoF(p)* is computed again, this time ignoring the values that are larger (in absolute value) than 2\**std_2_(p)*. Denote the result as *std_3_(p)*. This computation reduces the effect of outliers and transients on the computed threshold, which was set to *4*std_3_(p)*. For each representative pixel p, the values in *dFtoF(p)* which are higher than the *4*std_3_(p)* are considered to be part of a calcium transient (marked in green in Fig. 5B). Close (in time) calcium transient segments are unified by a morphological close operation (radius=2s) which closes small gaps between transients. A single detected transient that has more than one local maximum of *dFtoF* might be separated into several transients by the following steps. For each two consecutive local maxima of the same detected transient, which are at least 3 seconds apart, we check if the minimal *dFtoF* value between them is lower than the smaller local maxima by at least *2*std_3_* (significant “valley” between these two peaks). In case it is, this transient will be split into two transients at the minimal point between the two local maxima. Lastly, transients that overlap the first and last minute were ignored (as the computation of the baseline in these minutes is less accurate). Regions with fewer than 5 detected active cells were not included in the analysis. Since we saw that the calcium transients activity usually decreased during the recording session, we used the most active 2 minutes of each recording session for the analysis (i.e., the two minutes with the maximal number of detected calcium transients). Some of the recordings (usually the first recording per coverslip) had some spatial drift (slow movement of the coverslip) during the recording sessions, which were corrected by finding the shift between each frame and the first frame in the video. Shifts were computed by selecting the shift that maximizes the normalized cross-correlation with the first frame of the video. In such recordings, statistics were computed only for the region that is overlapped in all of the frames. When computing the number of active cells for such recordings, the number of active cells was multiplied by the ratio between the analyzed area and the full image area.

### Statistical analysis

Pairwise comparisons were computed using a non-parametric two-sided Wilcoxon rank sum test (Matlab’s ranksum function). P-values equal or smaller than 0.1, 0.05, 0.01, and 0.001 were marked by #, *, **,and ***, respectively. In addition, in the comparison of micturition and defecation rate, we used a two-way chi-square test to compare the number of zeros and non-zeros in WT vs Dup. FDR correction (Benjamini and Hochberg procedure) [34] was used in panels that include more than one pairwise comparison. The statistical data, including p-values (before and after FDR correction), and the data points for all of the statistical comparisons in this work are provided in Supplementary file 1.

## RESULTS

Since weaning, WT and Dup littermates were kept in mixed home-cages assigned to two conditions: Baicalin treatment or no treatment. In the Baicalin treatment cages, Baicalin was supplied via the drinking water (0.15 mg/ml) from the day of weaning (20-25 days) and until the end of the social behavior experiments, which were done at the age of 2-5 months (See Fig. 1A and Methods). In our analyses, we have considered four groups: WT with no treatment (WT), Dup with no treatment (Dup), Dup with Baicalin treatment (Dup-Bai), and WT with Baicalin treatment (WT-Bai).

### Social discrimination tests

Each subject mouse underwent three social discrimination tests, conducted on consecutive days. Each test comprised of a 15-minute habituation period with empty chambers in the arena, followed by a 5-minute trial period with two stimulus-containing chambers located at opposite corners of the arena (Fig. 1B-F and Methods). In the Social Preference (SP) test, a social stimulus (novel age- and sex-matched mouse) was positioned in one chamber while a Lego toy was positioned in the other chamber, located in the opposite side. In the Sex Preference (SxP), a male mouse was positioned in one chamber and a female mouse in the other. In the Emotional State Preference (ESP), a stressed mouse was positioned in one chamber and a naïve mouse in the other. A significantly higher time dedicated by the subject animals to investigate one of the stimuli over the other reflects a preference toward this stimulus. Male WT and Dup mice showed normal preference towards the social stimulus in SP test, the female stimulus in SxP test, and the stressed mouse stimulus in ESP test (Fig. 1G-I). Female WT and Dup mice showed similar behavior as males, with the exception of a preference towards the male stimulus during SxP exhibited by female Dup subject mice, which was not present for the WT female mice. (Fig. 1J-L). Baicalin treatment did not affect any of these preferences in either male or female subject mice (Fig. 1G-L), as also reflected by the Relative Discrimination Index (RDI) values (Supplementary Fig. 1A-C) with the exception of increased preference towards the male stimulus in female WT-Bai as compared to the female WT group (Supplementary Fig. 1D).

#### Baicalin ameliorates modified surface temperature in Dup male and female mice

The subject mice’s surface temperature was measured using infrared thermography (IRT) in the same manner as previously reported [8], at three body parts: eyes, center (body center), and tail-base during both habituation and trial periods of each test (Fig. 2A-B). When pooling together the results for the three social discrimination tests (SP, SxP, and ESP, see Supplementary Fig. 2,3 for each test), we found higher surface temperature in male Dup mice in all cases besides Center during the trial period (Fig. 2C-K). Baicalin treatment significantly reduced the temperature of Dup male mice in the tail-base and in a borderline significant manner in the eyes, for both the habituation and trial periods (Fig. 2C-K). A non-significant reduction of temperature in Baicalin-treated Dup male mice is also visible in the body center (Fig. 2D,G,J). For female Dup mice, we found a significantly colder tail-base temperature during the trial, as compared to WT females. Interestingly, Baicalin treatment increased the tail-base temperature during the trial period in Dup females. This suggests that the effect of Baicalin treatment is not just a cooling-down effect but rather reversing the influence of the Dup mutation on the surface temperature, as it reversed the higher temperature phenotype in male Dup eyes and tail-base, while also reversing the lower tail-base temperature phenotype in Dup females. Notably, in the WT group, Baicalin treatment increased the center (during habituation) and tail (during habituation and trial) temperature in males while decreasing the eyes temperature in females (Supplementary Fig. 3). Thus, the Baicalin effect on the surface temperature of WT mice is different and sometimes opposite (for example, in the males’ tail-base) to its effect in Dup mice, suggesting again a specific effect on Dup mice.

#### Baicalin ameliorates the higher defecation rate found in Dup male mice during the SP test

We used the DeePosit software and method [30] to track micturition and defecation rates during the trial period of the three behavioral tests (see examples in Fig. 3A,D). We found a higher defecation rate specifically during the SP trial in Dup males, but not females, compared to their WT littermates. Baicalin treatment reduced both the rate of defecation (borderline significance) and micturition (Fig. 3B,E) in Dup males, with no effect on WT littermates (Supplementary figure 5). Higher defecation rate in Dup male mice was also found in the ESP test with no significant effect to the Baicalin treatment (Supplementary Fig. 4A,C). No statistically significant effect of Baicalin treatment was found for Dup female mice (Fig. 3C,F, and Supplementary Fig. 4B,D).

Overall, our results so far in Dup mice suggest that Baicalin treatment alleviates most changes associated with *Gtf2i* duplication in the surface temperature and defecation activity during social interaction. We therefore decided to examine whether Baicalin treatment also affects the activity of neurons differentiated from 7Dup patient-derived iPSCs.

### Baicalin increases the excitatory post-synaptic currents (EPSC) rate in 7Dup patient-derived neurons while reducing the number of active cells in calcium imaging

Patient-derived cortical neurons were generated by reprogramming peripheral blood mononuclear cells taken from a 7Dup patient male child (7.5 years old at the time of taking the blood sample), as well as his father, as a control. Though we also report the measurements for the control line, we acknowledge that it is difficult to reach a conclusion by comparing a single non-isogenic control with a single patient line. Hence, we focus on the effect of the Baicalin treatment on the patient line.

Whole-cell patch clamp recordings (illustrated in Fig. 4A) were done at two time points (TPs): 33-38 (TP1) and 74-79 (TP2) days after the start of differentiation from NPC to neurons. Spontaneous action potentials recordings were done in current clamp mode and were analyzed by a custom Matlab-implemented peak detector (see Methods and Fig. 4B). No significant difference was found between treated and non-treated neurons in the frequency or amplitude of spontaneous action potentials (Fig. 4C-D). Evoked action potentials were recorded while injecting current steps into the patched cell (first 10 steps are illustrated in Fig. 4E-F). The mean number of action potentials per step was measured (Fig. 4G-H), but no significant difference was found between treated and non-treated cells (Fig. 4I).

Spontaneously occurring EPSCs were recorded in voltage clamp mode. Recordings were analyzed with a custom template-matching-based peak detector (see Methods, Fig. 4J and Supplementary Fig. 6). EPSC rate was higher in the Baicalin-treated cells in TP2 (Fig. 4K).

To check if the higher rate of EPSCs is related to higher neuronal connectivity or to a higher level of neuronal activity, we analyzed the activity of the cell population using calcium imaging, which captures the calcium activities of large groups of cells. 12-minute videos were captured from typically 10 regions in each well (first 10 minutes were analyzed). A custom automatic active cell detector was used to detect and segment active cells and to select a representative pixel for each cell. The most active two-minutes period of each region was used for analysis (see Methods, Fig. 5A, and Supplementary Fig. 7). Calcium transients for each representative pixel were analyzed using a custom peak detector (example of a peak detection results is illustrated in Fig. 5B). Lower number of active cells (during the most active 2 minutes of the recording) were found in the Baicalin treated 7Dup cells (Fig. 5C). However, the number of calcium transients per detected cell (during the most active 2 minutes of the recording) remained unchanged (Fig. 5D).

Thus, the Baicalin-treated 7Dup cells showed a decrease in the calcium activity per region, which, together with the higher EPSC rate, suggests a higher level of network connectivity rather than of neuronal activity. Altogether, these results suggest that Baicalin affects the synaptic activity of patient-derived 7Dup neurons in culture.

## DISCUSSION

The *GTF2I* gene is thought to be a main player in the 7Dup syndrome, caused by duplication of a 1.5-1.8 Mb segment in human chromosome 7. Dup mice, carrying a duplication of the *Gtf2i* gene, were previously used as a mouse model for 7Dup [8–11]. In this study, we explored the effect of Baicalin, a naturally occurring flavonoid commonly used in traditional Chinese medicine [12], on behavioral and physiological modifications exhibited by Dup mice during social discrimination tests, as well as its influence on the electrophysiological activity of 7Dup patient-derived neurons in culture.

LSD1 inhibition (LSD1i) and HDAC inhibition (HDACi) were previously suggested as potential treatments for 7Dup [11, 19, 20]. More specifically, one study [19] found that the mRNA level of BEND4, a transcription factor highly expressed in the brain and considered a potential risk gene for ASD, is downregulated in 7Dup patient-derived iPSCs and that LSD1i treatment rescues BEND4 mRNA levels in these cells. Another study [11] revealed deficits in sociability and in preference towards a novel social stimulus in Dup mice, and found that LSD1i treatment rescued this phenotype. A third study [20] showed that drugs with HDACi properties lowered GTF2I mRNA and protein levels in 7Dup patient-derived neurons. Therefore, we looked for molecules that exert LSD1i and HDACi properties, in order to explore their influence on behavioral and physiological deficits previously discovered by us in Dup mice [8].

The LSD1i and HDACi properties, as well as the HDAC downregulation activity of Baicalin were previously reported [23–27]. Specifically, one study [23] reported that Baicalin inhibits LSD1 with an IC_50_ of 3.01 μM and that Baicalin treatment increased the H3K4me2, a substrate of LSD1, in gastric cancer cell line MGC-803 without impact on LSD1 expression. Dilution of the Baicalin concentration resulted in recovery of the LSD1 activity in these cells, suggesting that the Baicailn effect is reversible. Another study [27] found that Baicalin inhibits HDAC of class I, IIa, IIb in bovine cardiac tissue with IC_50_ of 52 μM, 67 μM, 85 μM, respectively. A third study [25] found that Baicalin treatment (Intrathecal injection) downregulated HDAC1 expression in the spinal cords of a neuropathic pain rat model. In addition, they found increased H3 acetylation (which might be related to LSD1i activity) in the spinal cord. However, another study [35] reported that Baicalin treatment restored downregulated levels of HDAC2 protein in cigarette smoke-induced inflammatory mouse model and cell model, suggesting that Baicalin does not always inhibit HDAC, but rather has a more complex mechanism of action. When administered orally, the effect of Baicalin may also be mediated by its metabolites, which are generated by the gut microbiome [36]. These metabolites include Baicalein, the aglycone derivative of Baicalin, which was also reported to have an HDACi activity [27]. Altogether, these studies suggest that Baicalin administration may serve as a treatment inducing LSD1i and HDACi activity. Therefore, here we treated Dup mice with Baicalin and examined the effect of this treatment on the behavioral and physiological deficits of Dup mice.

Our main observation is that Baicalin treatment normalized the modified surface temperature exhibited by Dup mice during social discrimination tests. Baicalin’s effect on the mice’s surface temperature has several potential mechanisms. In our previous work [8], we showed that *Gtf2i* dosage correlates with mice surface temperature in both *Gtf2i*^+/dup^ and *Gtf2i*^+/−^ mice. These results suggested that inhibition of *Gtf2i* expression or activity in Dup mice would normalize their surface temperature. Interestingly, Baicalin was found to have an antipyretic effect in a rabbit fever model [37] where it also reduced the expression of the inflammatory cytokine TNF-α, the level of which is also associated with thermoregulation [38]. These results are consistent with previous reports on Baicalin’s anti-inflammatory properties [16]. Interestingly, while Baicalin reduced the surface temperature in Dup male mice, thus normalizing their phenotype, it also increased the tail temperature in Dup female mice, also ameliorating their phenotype in comparison to WT female mice. The colder tail temperature in Dup females might be related to anxiety-induced vasoconstriction, as was also found in [8] in response to stress, and hence might be ameliorated by the anxiolytic effect of Baicalin. In any case, the fact that Baicalin treatment normalized the opposite modifications in surface temperature exhibited by male and female Dup mice excludes the possibility that its effect is due to a general warming or cooling effect. Instead, it supports our hypothesis that Baicalin treatment is ameliorating the effects of *Gtf2i* duplication in both male and female Dup mice.

Accordingly, Baicalin treatment also normalized the increased defecation activity of Dup male mice during the Trial period of the SP test. In a previous work, we found opposite changes in defecation rate during the SP test in male mice with duplication and deletion of the Gtf2i gene, suggesting that Gtf2i dosage dictates this behavioral variable. The fact that this dependence was found only for the SP test may be related to the fact that this is the first test conducted, hence it may be associated with a higher level of anxiety. In any case, the fact that no change in defecation rate was observed in the SxP test proves that it is not related to metabolic changes in Dup mice, but rather to their behavior. It should be noted, however, that the effect of Baicalin treatment on defecation rate was limited to the SP test and did not repeat itself on the elevated rate exhibited by Dup male mice in the ESP test, hence seems to be less consistent than the effect of Baicalin on surface temperature, which was apparent, at least as a trend, across all tests.

We found that Baicalin treatment (5ug/ml, which is 11.2uM) of 7Dup patient-derived neurons increased the frequency of EPSCs measured with whole cell patch clamp. However, calcium imaging of the neuronal cultures showed a reduced number of active cells per region and no change in the number of calcium transients per active cell. These results imply that the increased frequency of EPSCs might be related to higher synaptic activity in the Baicalin-treated cell cultures, in accordance with the findings of a previous study [39], which reported increased neuronal outgrowth in Baicalin-treated NT2-N neuronal-like cells (human-derived). Interestingly, Deurloo et al [10] found reduced neurite complexity (reduced neurite length and reduced number of axon branches) in mice with duplicated *Gtf2i* and the opposite in mice with *Gtf2i* deletion, supporting a potential role of Baicalin as a potential treatment for ameliorating this reduced neuronal complexity.

In conclusion, Baicalin treatment ameliorates behavioral and physiological modifications in mice with duplicated *Gtf2i* gene and affects synaptic activity in 7Dup patient-derived cultured neurons, supporting Baicalin as a potential candidate for treating 7Dup patients.

### Limitations

In the animal research, we used three batches of experiments carried out over a long period (∼2 years), including different seasons and during different hours of the day. Only the last batch of these experiments included the Baicalin treatment. We started the treatment at a relatively young age, hence we cannot determine if Baicalin treatment starting at older ages would be effective. Also, we did not measure the surface temperature of Dup animals in their home cages, hence we do not know whether their modified surface temperature is constant or associated with the behavioral tests. In the patient-derived neurons experiment, we used a single patient and a single non-isogenic control. Hence, we cannot draw conclusions regarding the difference between 7Dup and Control-derived neurons. We therefore limit our conclusions to the effect of Baicalin treatment on the 7Dup cell line that we measured.

## ACKNOLEDGEMENTS

We thank Prof. Lucy Osborne and Prof. Giuseppe Testa for providing the *Gtf2i*^+/dup^ mice. We thank Yaniv Goldstein, Janet Tabakova, Wjdan Awaisy, Shorook Amara, and Ori Yacobi for their help in annotating the videos and Sara Sheikh for drawing the experiment setup illustration. We thank Haya Kaabeya for performing the differentiation process from iPSCs to NPCs.

## DATA AVAILABILITY

The dataset would be made available from the corresponding author upon reasonable request.

## CODE AVAILABILITY

The code used for analysis would be made available in a suitable public repository from the corresponding author upon acceptance of the manuscript.

## AUTHOR CONTRIBUTIONS

D.P: Data curation, Formal analysis, Investigation, Methodology, Writing – original draft, Algorithms and software, Visualization, Cell work.

S.N: Data curation, Project administration, Resources, Algorithms and software, Supervision.

N.R: Data curation, Investigation.

T.S: Data curation, Investigation.

S.S: Funding acquisition, Supervision, Writing – review and editing.

S.W: Funding acquisition, Supervision, Writing – review and editing.

