## Supplementary Figures for "Baicalin ameliorates behavioral and physiological deficits in *Gtf2i*-duplication mice and modulates synaptic activity in 7q11.23 Duplication Syndrome patient-derived neurons"

SUPPLEMENTAL FIGURES

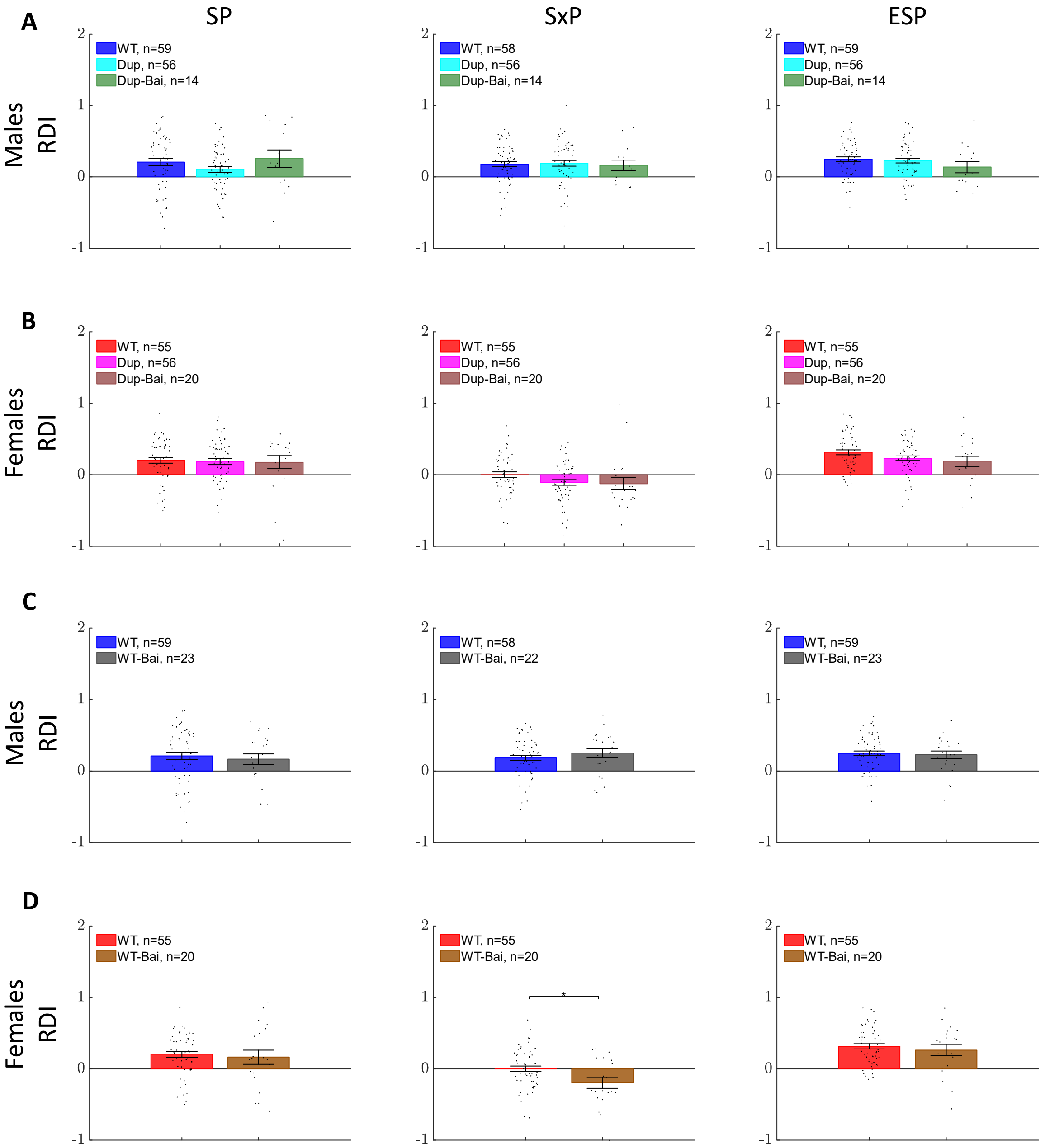


***Supplemental Figure 1: RDI at SP, SxP, and ESP****. Mean RDI values ±SEM for males (**A,C**) and Females (**B,D**) during SP, SxP and ESP tests. Positive values indicate preference towards stimulus 1, which is the social in the SP test, the female in the SxP test, and the stressed mouse in ESP. Negative values indicate a preference towards stimulus 2. Pairwise comparisons between WT and Dup, between Dup and Dup-Bai, and between WT and WT-Bai were done using Wilcoxon rank sum test. Panels in (**A-B**) were FDR corrected (2 comparisons per panel).*

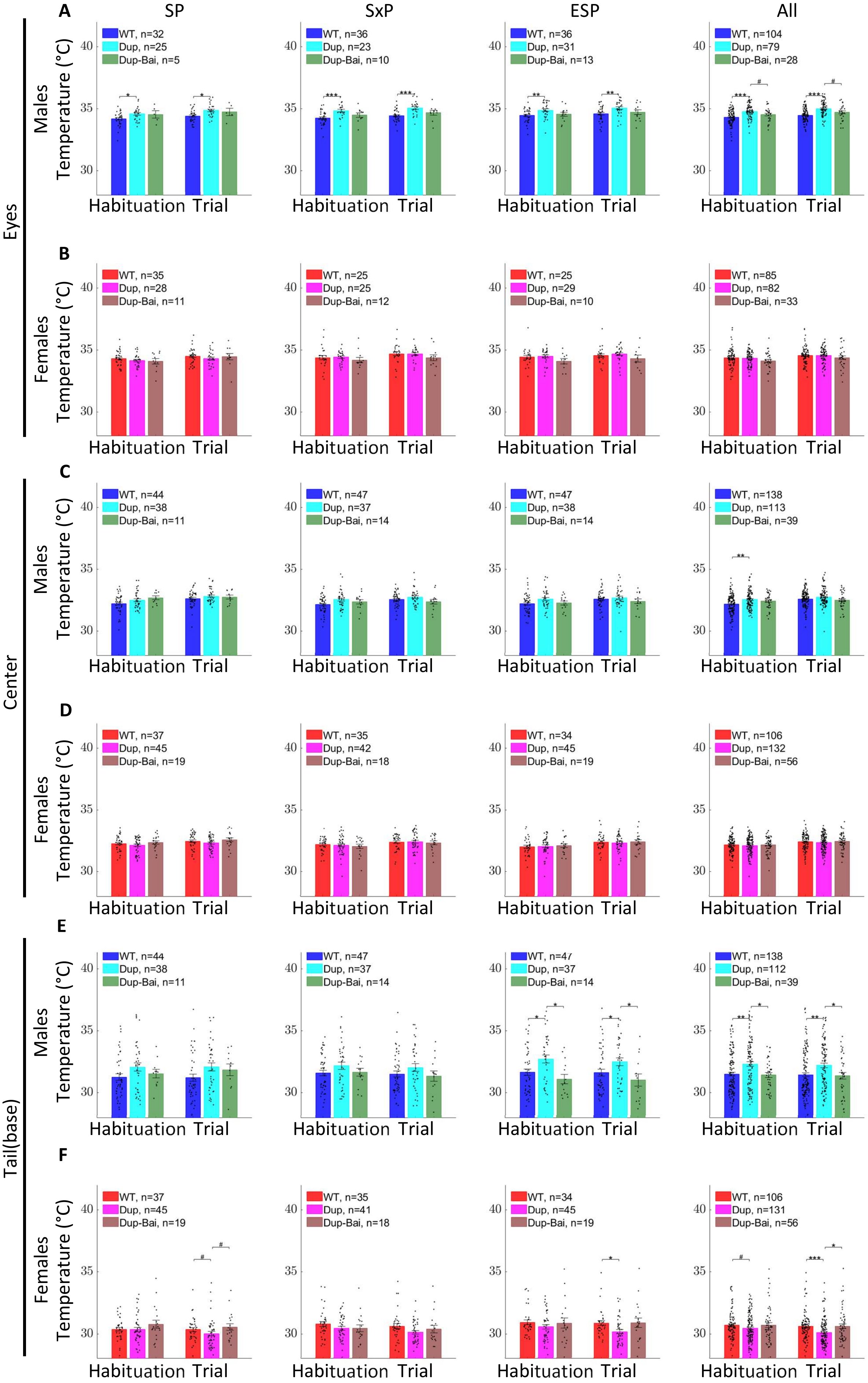

A
SP
SxP
ESP
All
Eyes
Males
Temperature (°C)
WT, n=32
Dup, n=25
Dup-Bai, n=5
WT, n=36
Dup, n=23
Dup-Bai, n=10
WT, n=36
Dup, n=31
Dup-Bai, n=13
WT, n=104
Dup, n=79
Dup-Bai, n=28
Habituation Trial
B
Females
WT, n=35
Dup, n=28
Dup-Bai, n=11
WT, n=25
Dup, n=25
Dup-Bai, n=12
WT, n=25
Dup, n=29
Dup-Bai, n=10
WT, n=85
Dup, n=82
Dup-Bai, n=33
C
Center
WT, n=44
Dup, n=38
Dup-Bai, n=11
WT, n=47
Dup, n=37
Dup-Bai, n=14
WT, n=47
Dup, n=38
Dup-Bai, n=14
WT, n=138
Dup, n=113
Dup-Bai, n=39
D
WT, n=37
Dup, n=45
Dup-Bai, n=19
WT, n=35
Dup, n=42
Dup-Bai, n=18
WT, n=34
Dup, n=45
Dup-Bai, n=19
WT, n=106
Dup, n=132
Dup-Bai, n=56
E
Tail(base)
WT, n=44
Dup, n=38
Dup-Bai, n=11
WT, n=47
Dup, n=37
Dup-Bai, n=14
WT, n=47
Dup, n=37
Dup-Bai, n=14
WT, n=138
Dup, n=112
Dup-Bai, n=39
F
WT, n=37
Dup, n=45
Dup-Bai, n=19
WT, n=35
Dup, n=41
Dup-Bai, n=18
WT, n=34
Dup, n=45
Dup-Bai, n=19
WT, n=106
Dup, n=131
Dup-Bai, n=56
30
35
40

***Supplemental Figure 2: Mice surface temperature in SP, SxP, and ESP.*** *Mean temperature ±SEM of the Eyes (**A-B**), Center (**C-D**), and Tail(base) (**E-F**) for male WT vs Dup and Dup vs Dup-Bai (**A,C,E**) and the same for females in (**B,D,F**), during habituation and trial periods of SP, SxP, ESP tests and during All tests pooled together. Pairwise comparisons were done using Wilcoxon rank sum test and were FDR corrected (4 comparisons per panel).*

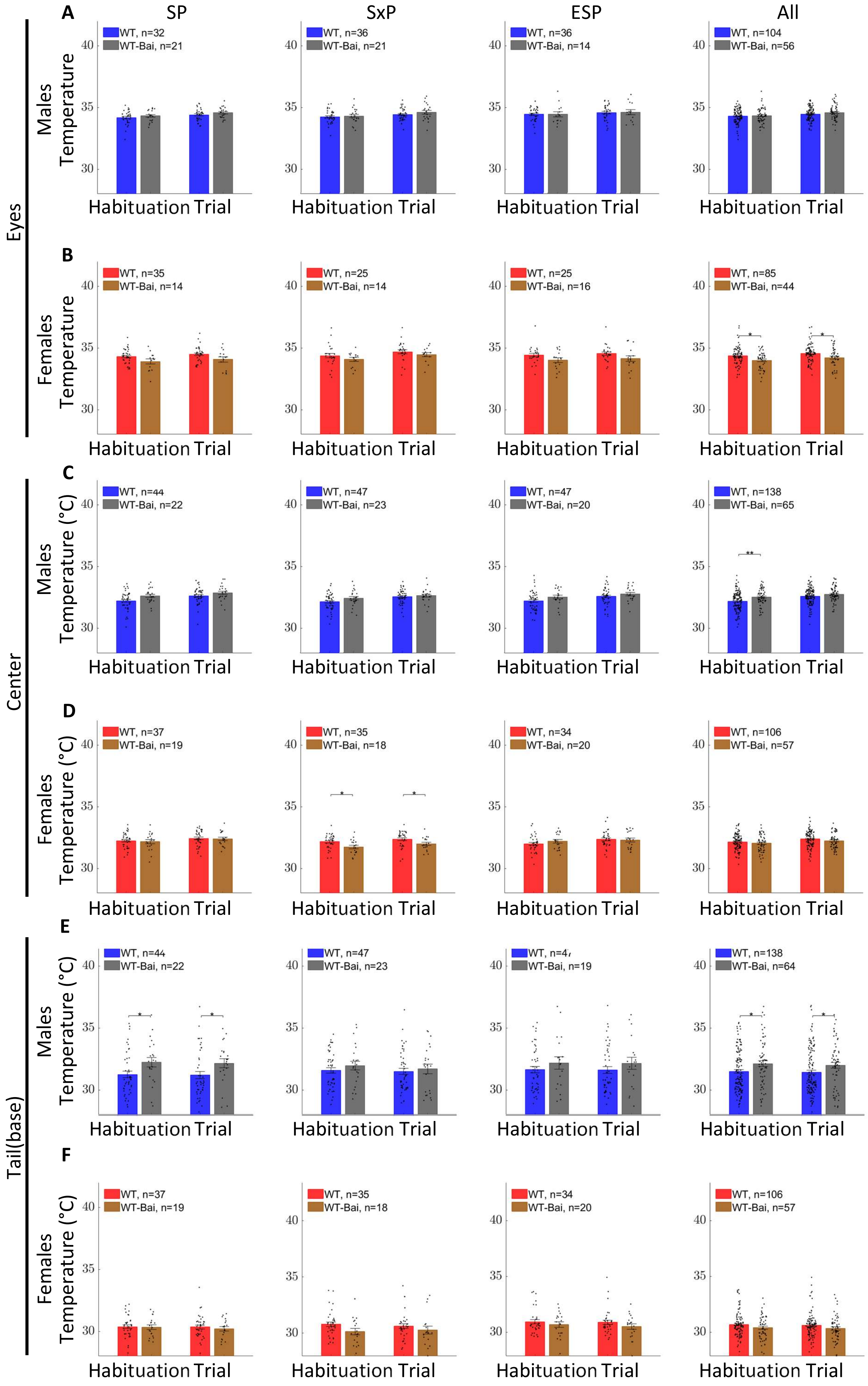

A
SP
SxP
ESP
All
Eyes
Males
Temperature
WT, n=32
WT-Bai, n=21
WT, n=36
WT-Bai, n=21
WT, n=36
WT-Bai, n=14
WT, n=104
WT-Bai, n=56
Habituation Trial
B
Females
Temperature
WT, n=35
WT-Bai, n=14
WT, n=25
WT-Bai, n=14
WT, n=25
WT-Bai, n=16
WT, n=85
WT-Bai, n=44
Habituation Trial
C
Center
Males
Temperature (°C)
WT, n=44
WT-Bai, n=22
WT, n=47
WT-Bai, n=23
WT, n=47
WT-Bai, n=20
WT, n=138
WT-Bai, n=65
Habituation Trial
D
Females
Temperature (°C)
WT, n=37
WT-Bai, n=19
WT, n=35
WT-Bai, n=18
WT, n=34
WT-Bai, n=20
WT, n=106
WT-Bai, n=57
Habituation Trial
E
Tail(base)
Males
Temperature (°C)
WT, n=44
WT-Bai, n=22
WT, n=47
WT-Bai, n=23
WT-Bai, n=19
WT, n=138
WT-Bai, n=64
Habituation Trial
F
Females
Temperature (°C)
WT, n=37
WT-Bai, n=19
WT, n=35
WT-Bai, n=18
WT, n=34
WT-Bai, n=20
WT, n=106
WT-Bai, n=57
Habituation Trial
30
35
40

***Supplemental Figure 3: Mice surface temperature in SP, SxP, and ESP for WT vs WT-Bai.*** *Mean temperature ±SEM of the Eyes (**A-B**), Center (**C-D**), and Tail(base) (**E-F**) for male WT vs WT-Bai (**A,C,E**) and the same for females in (**B,D,F**), during habituation and trial periods of SP, SxP, ESP tests, and during All tests pooled together. Pairwise comparisons were done using Wilcoxon rank sum test and were FDR corrected (2 comparisons per panel).*

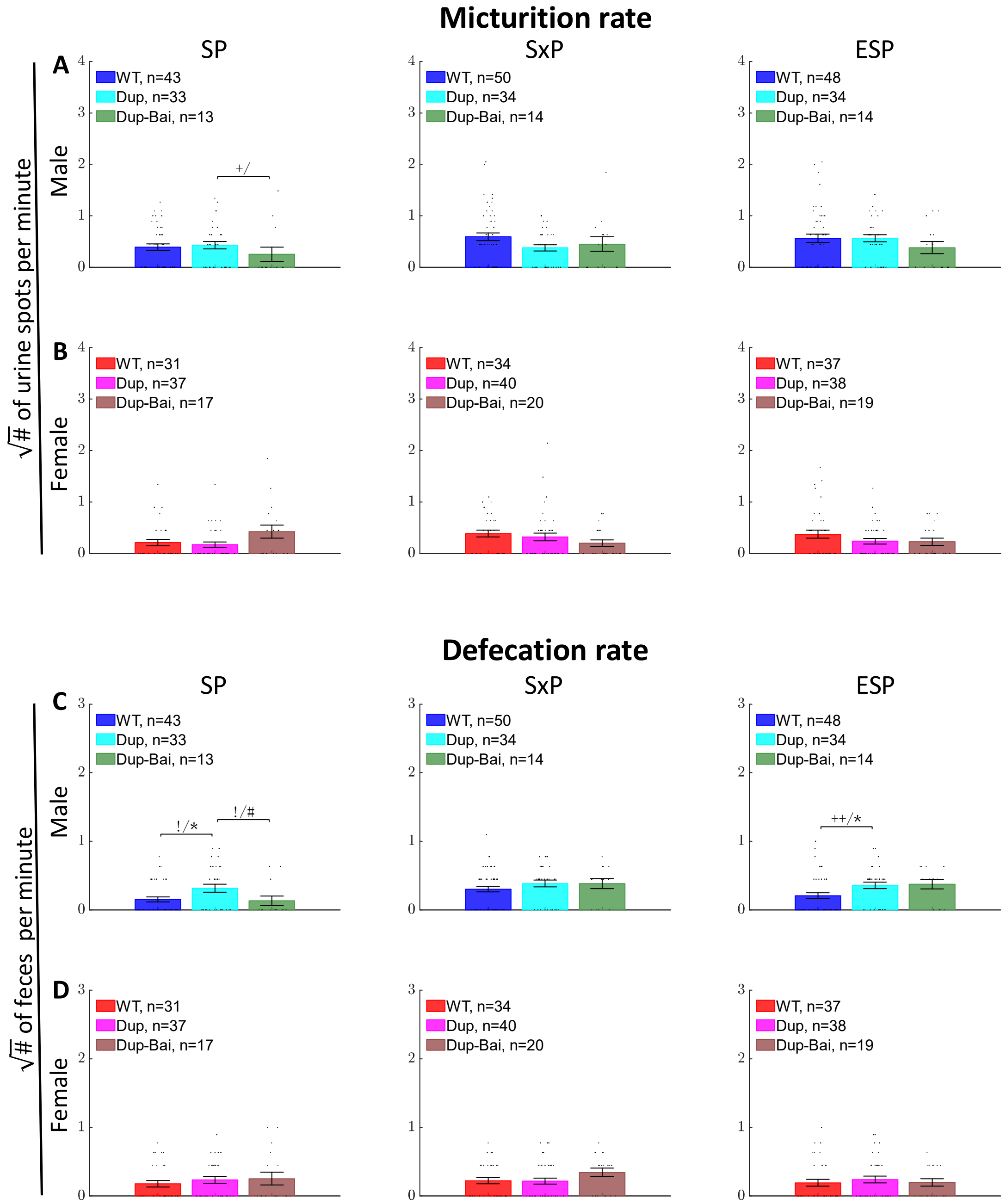


***Supplemental Figure 4: Micturition and defecation rate during SP, SxP and ESP.*** *Mean micturition rate ±SEM (**A, B**) and defecation rate (**C, D**) for WT vs Dup and Dup vs Dup-Bai for male (**A, C**) and female (**B, D**) mice during trial period of SP, SxP and ESP tests. FDR-corrected (2 comparisons per panel) pairwise comparisons using two-sided Wilcoxon ranksum test were marked with #,* for p<0.1 and p<0.05, respectively. Pairwise comparisons using two way Chi-square test were used to compare the distribution of zeros between each two compared groups. Significance in the Chi-square test (FDR-corrected, 2 comparisons per panel) was marked with !,+ for p<0.1 and p<0.05, respectively.*

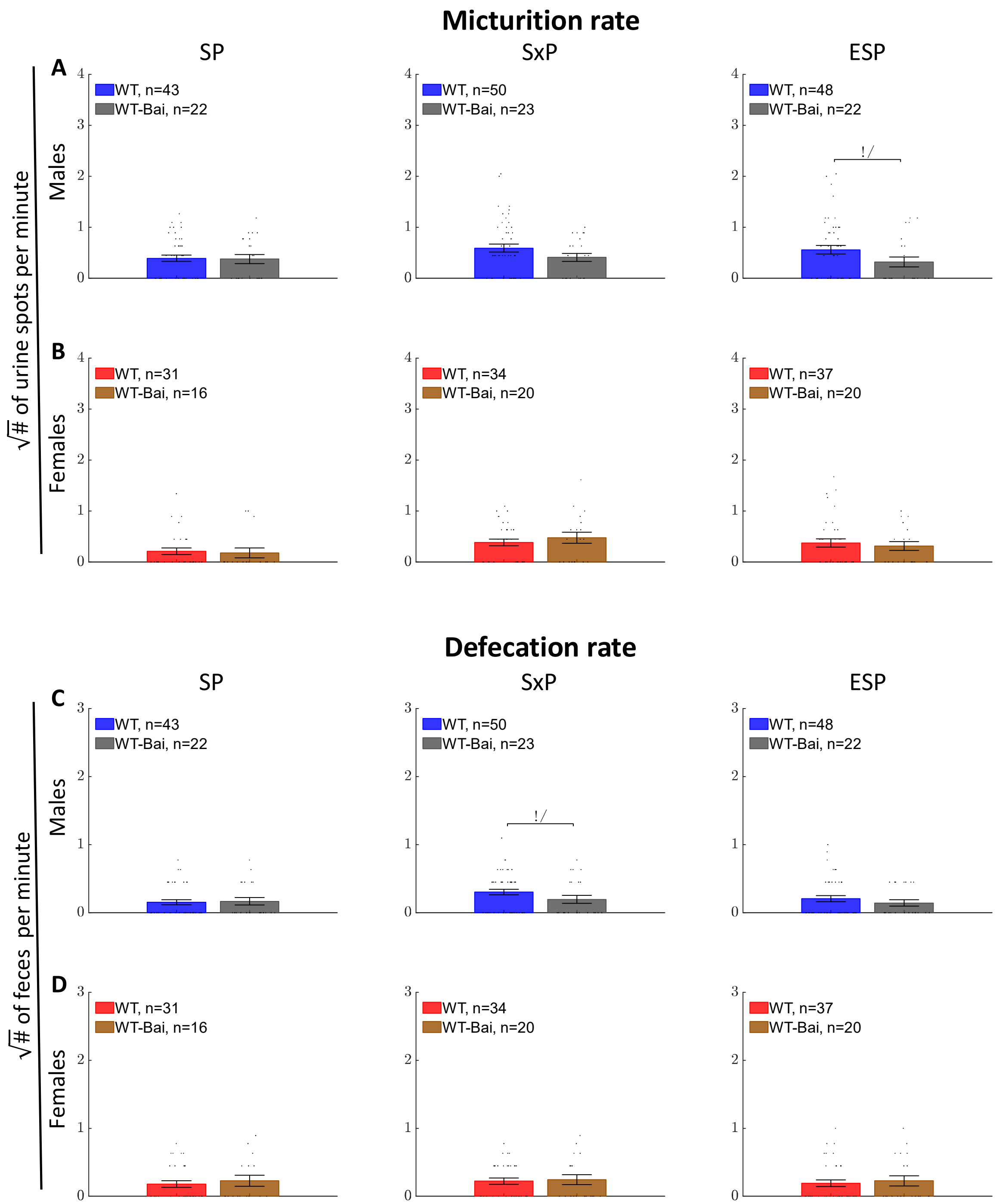


*Supplemental Figure 5:* ***Micturition and defecation rate during SP, SxP, and ESP for WT vs WT-Bai.*** *Mean ±SEM micturition rate (**A, B**) and defecation rate (**C, D**) for male WT vs WT-Bai (**A, C**) and female (**B, D**) mice during the trial period of SP, SxP and ESP tests. Pairwise comparisons using the two-sided Wilcoxon rank-sum test were not statistically significant. Pairwise comparisons using a two-way Chi-square test were used to compare the distribution of zeros between each pair of compared groups and were marked with ! for p<0.1.*

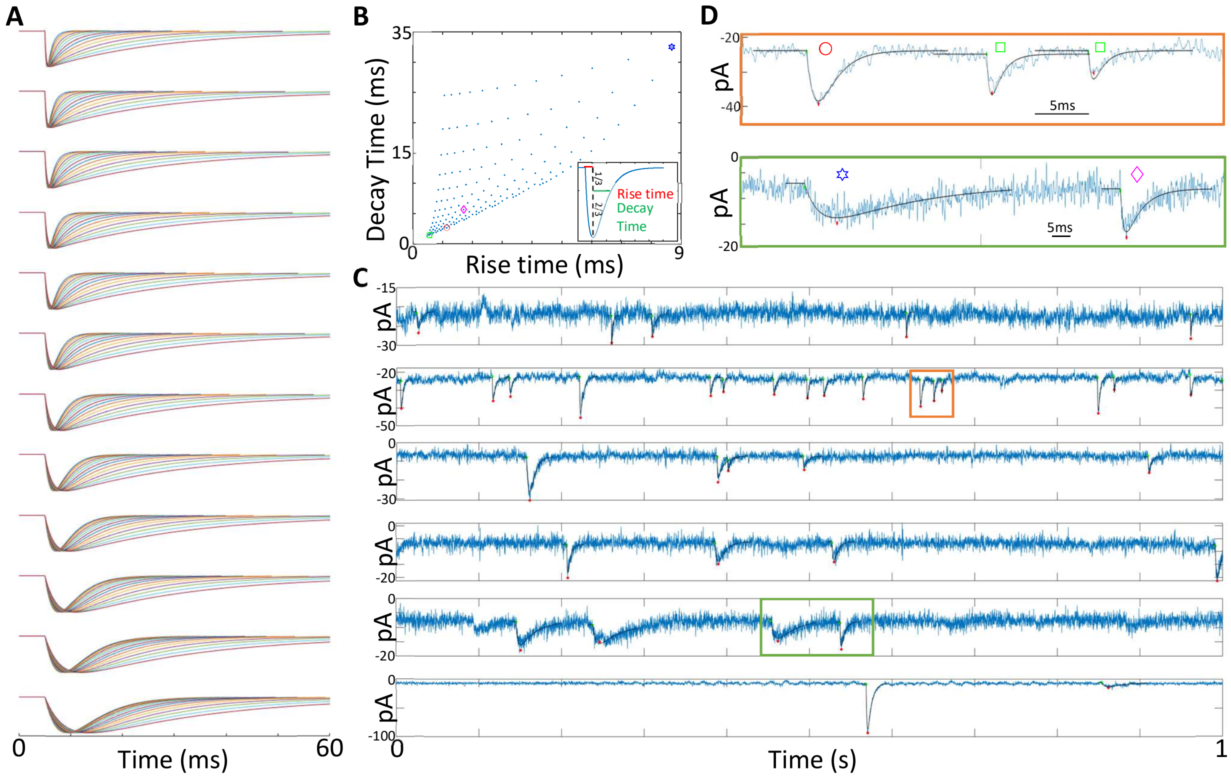


***Supplementary Figure 6: Automatic EPSCs detection.*** *(**A**) A set of 168 synthetic EPSCs profiles were generated and used for template matching as part of the EPSCs detection algorithm. (**B**) The 168 synthetic profiles shown in (**A**) have the rise times and decay times (recovery of 2/3 of the amplitude) shown in (**B**). (**C**) Example of automatic EPSCs detection results. A green dot marks the onset of an EPSC, and a red dot marks the minimal value. The matched synthetic profile is overlayed in black. In the bottom row, a detection with a rise amplitude greater than 30 pA was considered to be an action potential and not an EPSC. (**D**) Magnification of two regions in (**C**). The symbol above each EPSC in (**D**) matches the profile with the same symbol in (**B**).*

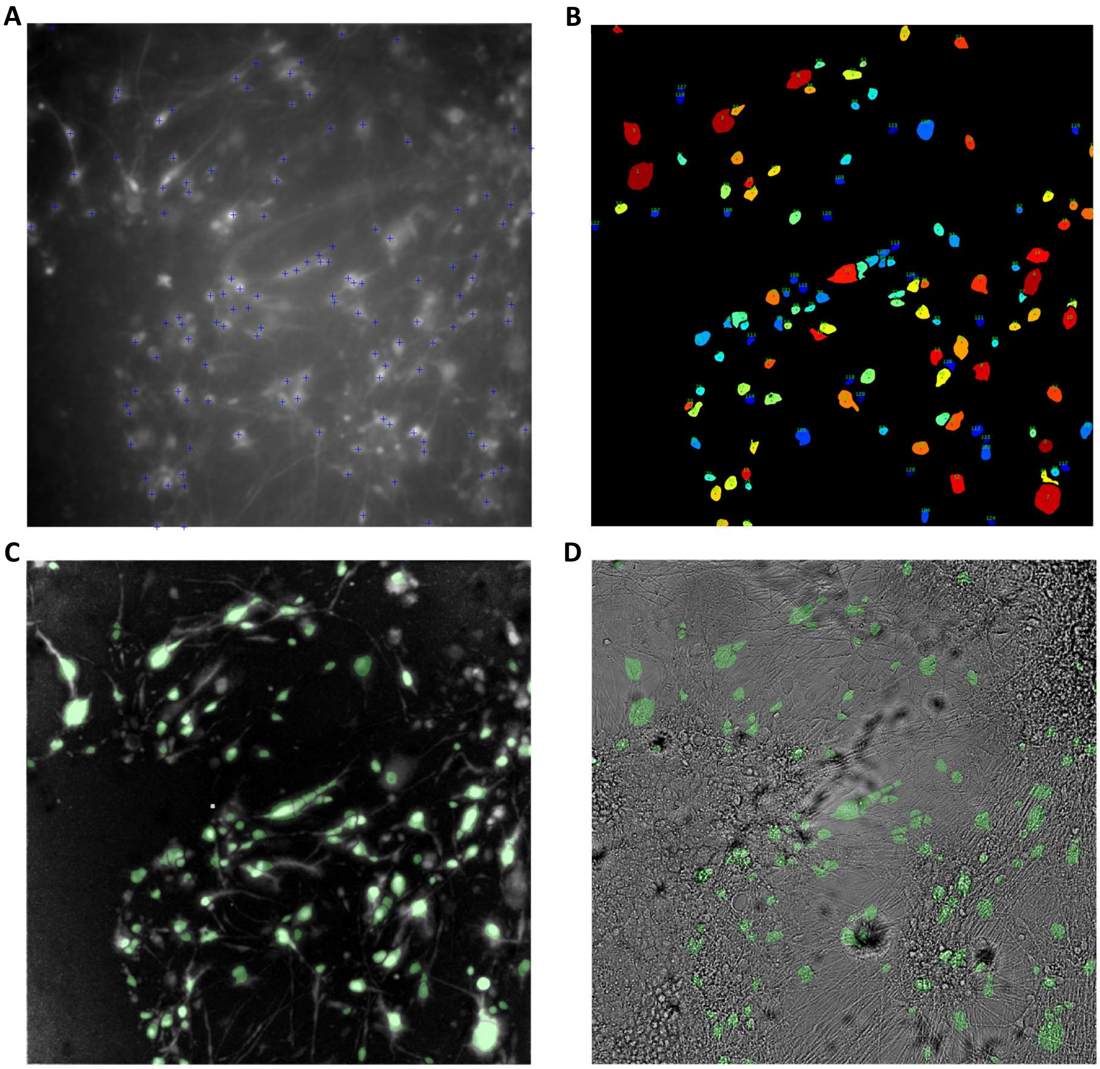


***Supplementary Figure 7: Automatic detection of active cells in calcium imaging video.*** *(**A**) Mean fluorescence image (of 10 minutes recording) overlayed by the detected units' representative pixels. (**B**) Detected active units segments (**C**) Active units segments are overlayed in green on the calcium activity map (maxHtoL image ,see Methods). (**D**) Active units segments are overlayed in green on the brightfield image.*
